# Genome-wide identification of *Pseudomonas syringae* genes required for competitive fitness during colonization of the leaf surface and apoplast

**DOI:** 10.1101/637496

**Authors:** Tyler C. Helmann, Adam M. Deutschbauer, Steven E. Lindow

## Abstract

The foliar plant pathogen *Pseudomonas syringae* can establish large epiphytic populations on leaf surfaces before infection. However, the bacterial genes that contribute to these lifestyles have not been completely defined. The fitness contributions of most genes in *P. syringae* pv. *syringae* B728a were determined by genome-wide fitness profiling with a randomly barcoded transposon mutant library that was grown on the leaf surface and in the apoplast of the susceptible plant *Phaseolus vulgaris*. Genes within the functional categories of amino acid and polysaccharide (including alginate) biosynthesis contributed most to fitness both on the leaf surface (epiphytic) or in the leaf interior (apoplast), while genes in the type III secretion system and syringomycin synthesis were primarily important in the apoplast. Numerous other genes that had not been previously associated with *in planta* growth were also required for maximum epiphytic or apoplastic fitness. Many hypothetical proteins and uncategorized glycosyltransferases were also required for maximum competitive fitness in and on leaves. For most genes, no relationship was seen between fitness *in planta* and either the magnitude of their expression *in planta* or degree of induction *in planta* compared to *in vitro* conditions measured in other studies. A lack of association of gene expression and fitness has important implications for the interpretation of transcriptional information and our broad understanding of plant-microbe interactions.

**Significance Statement:** Many plant pathogenic bacteria can extensively colonize leaf surfaces before entry and multiplication within the leaf to cause disease. While these habitats presumably require distinct adaptations, the genes required in these habitats and how they would differ was unknown. Using a genome-wide library of barcoded insertional mutants in the plant pathogen *Pseudomonas syringae*, we ascertained the common and unique genes required to colonize these habitats. A lack of association between gene expression and contribution to fitness suggests that many genes that are highly expressed or induced *in planta* are dispensable or redundant. As a model bacterium for plant pathogenesis and colonization, our comprehensive genetic dataset allows us to better understand the traits needed for association with leaves.

## Introduction

Many plant pathogenic bacteria are capable of extensive colonization of leaf surfaces before their entry and multiplication within the leaf. As such, epiphytic (leaf surface) populations on asymptomatic plants are considered a reservoir of inoculum, that under the appropriate conditions can lead to infection. In many cases, the likelihood of disease can be predicted from the epiphytic population size of the pathogen several weeks before infection occurs (1). The ability to form large epiphytic populations therefore is a measure of success for such a pathogen, and factors that determine its ability to grow on leaves would be considered fitness factors. After entry into the apoplast, bacterial numbers often increase greatly and disease is associated with those sites in which large internal population sizes have been achieved (2). The ability to grow within the apoplast of plants is thus also a measure of its fitness. In addition to plant pathogens, a diversity of other bacteria and fungi typically colonize the surface of aboveground plant parts. Such commensal bacteria and fungi are, however, typically limited to epiphytic growth with only very small numbers of such taxa found within plant tissue as endophytes (3). It is presumed that the growth of epiphytic bacteria is supported by their consumption of a variety of carbon and nitrogen-containing compounds that transit from the interior of the plant to the leaf surface (4, 5). A variety of mono- and disaccharides are thought to constitute the majority of the carbon containing compounds on leaf surfaces, with smaller amounts of other sugars, organic acids, and amino acids also present (6, 7). The absolute amount of such nutrients on leaves is generally low, and the growth of epiphytic bacteria is typically carbon-limited (6). Furthermore, the abundance of such nutrient sources on plants is spatially heterogeneous (4, 6). Because of the apparent chemical complexity, and spatially heterogeneous chemical and physical features of leaves, those traits needed for epiphytic fitness remain largely uncharacterized (8). Only limited descriptions of the chemical and physical environment found within the apoplast of plants have appeared (9). While many of the nutrient resources on the surface are apparently also present in the apoplast, the chemical and physical environment there is largely unknown. Water availability apparently limits intercellular growth (10), and the ability of pathogens to induce plants to release water into the apoplast may be a major feature required for exploitation of this habitat (11, 12). While the apoplast provides bacterial cells protection from environmental stresses on the leaf surface, they are in intimate proximity to living plant cells, and thus subject to inhibitory responses by the plant mediated by the innate immune system (3, 7). Taken together, it is clear that a large repertoire of traits beyond resource acquisition, such as motility, habitat modification, and various interactions with the plant may be needed by a plant pathogenic bacterium to exploit both the leaf surface and the leaf interior. Unfortunately, very few such fitness traits beyond those associated with interactions with the plant immune system have been identified.

*Pseudomonas syringae* is a plant pathogenic bacterial species that includes strains pathogenic on a wide variety of different plant species (13). Most strains have a prominent epiphytic phase, especially on the plant species for which they can also cause infection. Strains of *P. syringae* are commonly found as epiphytes on a variety of both host-and non-host plants, both in agricultural systems as well as native plant communities (14, 15). Many strains are capable of catalyzing ice formation, and because they can be found in rainfall, pristine snow, as well as in water sources around the world, are thought to play an important role in the water cycle by initiating ice formation central to the precipitation process (15). Such a connection to precipitation may also serve as a vehicle for long-distance dispersal as well as a mechanism for migration to plants after dispersal (16).

The model strain *P. syringae* pv. *syringae* B728a (B728a) is a strong epiphytic colonizer that was originally isolated from green bean (*Phaseolus vulgaris*), and is capable of causing brown spot disease (17). It is the best-studied member within *P. syringae* phylogroup II. This monophyletic clade contains strains that are overrepresented in environmental samples and are generally better epiphytes than members of other clades (18, 19). In addition, phylogroup II contains many strains with broad host ranges (19). Strain B728a is also pathogenic on *Nicotiana benthamiana* (20) and pepper (*Capsicum annuum*) (21). Like other ice nucleation active strains of *P. syringae*, this strain contributes to frost damage in frost sensitive plant species by limiting their ability to supercool and avoid damaging ice formation (22). Strain B728a is also a model for phytotoxin production, and contains a much smaller type III effector repertoire than many other strains such as *P. syringae* pv. *tomato* strain DC3000 (23, 24). As such, it has been hypothesized that this lower type III effector repertoire belies an increased reliance on broad-spectrum toxins as well as ice nucleation ability that contributes to its broad host range and more general environmental distribution (25). This robust epiphytic colonizer and ubiquitous plant pathogen is thus a useful model to examine traits needed for bacterial success in diverse environments.

The genes that are putatively the most ecologically relevant to the success of bacteria on plants have typically been identified on the basis of their transcriptional induction or expression in a given habitat (26). Measurements of gene expression, directly via microarray or RNAseq, or indirectly through reporter genes or *in vivo* expression technology (IVET), have been used in *Pseudomonas syringae* to identify genes that have host-responsive expression patterns (27–31). Validation of the role of genes identified by this method however usually involves targeted disruption of such genes individually with subsequent assessment of changes in behavior. This is a laborious procedure that cannot be readily applied to the genome as a whole. The high variability of the population size of a given strain after inoculation makes the comparison of population sizes between mutants and parental strains difficult. Large numbers of replicate samples are required to distinguish differential growth of such strains unless they differ greatly (32). Moreover, transcriptional studies may be of limited use in identifying host-colonization genes due to the lack of correlation between gene expression and contribution of those genes to fitness that is often observed *in vitro* (33). Examples of such a lack of correspondence of gene expression and fitness contribution in some *Pseudomonas* species on hosts have appeared (34, 35) but it is unclear how prevalent such a lack of connection might be.

To ascertain the roles of individual genes in *P. syringae* during epiphytic and apoplastic colonization on a genome-wide scale, we utilized a highly parallel transposon-based genomic screen. A variety of techniques taking advantage of high-throughput sequencing have been used to identify genes contributing to host colonization (36, 37). For example, genomic comparisons can reveal differential gene abundance in strains and genes that are under putative positive selection, suggesting that they might be contributing to host-specific fitness (38–40). However, confirmation of the role of such genes typically requires laborious mutation analysis as discussed above. Random mutagenesis techniques can enable genome-wide, gene-specific fitness contributions to be measured in a given habitat. One such strategy employs transposon sequencing (TnSeq) wherein the relative proportion of a transposon mutant in a given gene within a mixture of such mutants is assessed both before and after the strain mixture experiences a given condition. The number and genome location of the mutants in the mixture is determined by determining the sequences adjacent to the transposon by high throughput sequencing in each experiment (41). Recently developed random-barcoded transposon sequencing (RB-TnSeq) (42) is a modification of TnSeq that enables a transposon library to be used more easily for multiple assays since tagged transposons are used in mutagenesis and need be mapped only once. Eliminating the need to re-map transposon insertions in each experiment dramatically reduces the effort to carry out fitness screens in multiple conditions by using a single library of mutants. In a transposon mutant pool where each transposon is linked to a unique 20-nucleotide barcode, insertion mutant fitness is calculated through amplicon sequencing of the barcode regions to calculate the relative abundance of a given strain. Change in barcode relative abundance over time is used as a proxy for strain fitness within the population. RB-TnSeq was recently used to identify genes required by *P. simiae* for its invasion of *Arabidopsis thaliana* roots (43). In this study, we used RB-TnSeq to identify genes in *P. syringae* needed for its colonization of both the surface and interior habitats of bean. Since the stimulon for these two habitats had previously been determined (15), we also addressed the extent to which transcriptional changes in gene expression were predictive of the fitness contributions of these same genes.

## Results

### Adapting RB-TnSeq for an epiphyte and foliar pathogen

In order to screen for genes in *P. syringae* strain B728a contributing to host colonization, we generated a randomly DNA barcoded *mariner* transposon library using the *Escherichia coli* donor library created by Wetmore *et al* (42). The sequenced B728a mutant library consisted of 281,417 strains with insertions that map to the B728a genome, each containing a unique DNA barcode. Computationally removing insertional mutants outside the central 10 – 90% of coding region of a given gene resulted in 169,826 genic strains for analysis, with a median of 21 insertions per gene. The number of usable insertional mutants for each gene was correlated with the number of TA dinucleotide sites within each coding region (Pearson correlation coefficient r = 0.72) (Fig. S1). We analyzed fitness contributions for 4,296 of 5,137 (84%) protein-coding genes that harbored sufficient insertions for analysis.

The rich medium King’s B (KB) was used for library recovery prior to plant inoculations, so overnight growth in this condition was used as the control against which growth of the mutants on the leaf surface and in the apoplast (Fig. 1) was compared. All experiments analyzed herein passed quality control metrics that were previously established for *in vitro* studies in (42). The requirements for a successful experiment include ≥ 50 median reads per gene and consistency in the calculated fitness estimate obtained from mutants with insertions in the 3’ and 5’ half of a gene (42).

**Fig. 1.**
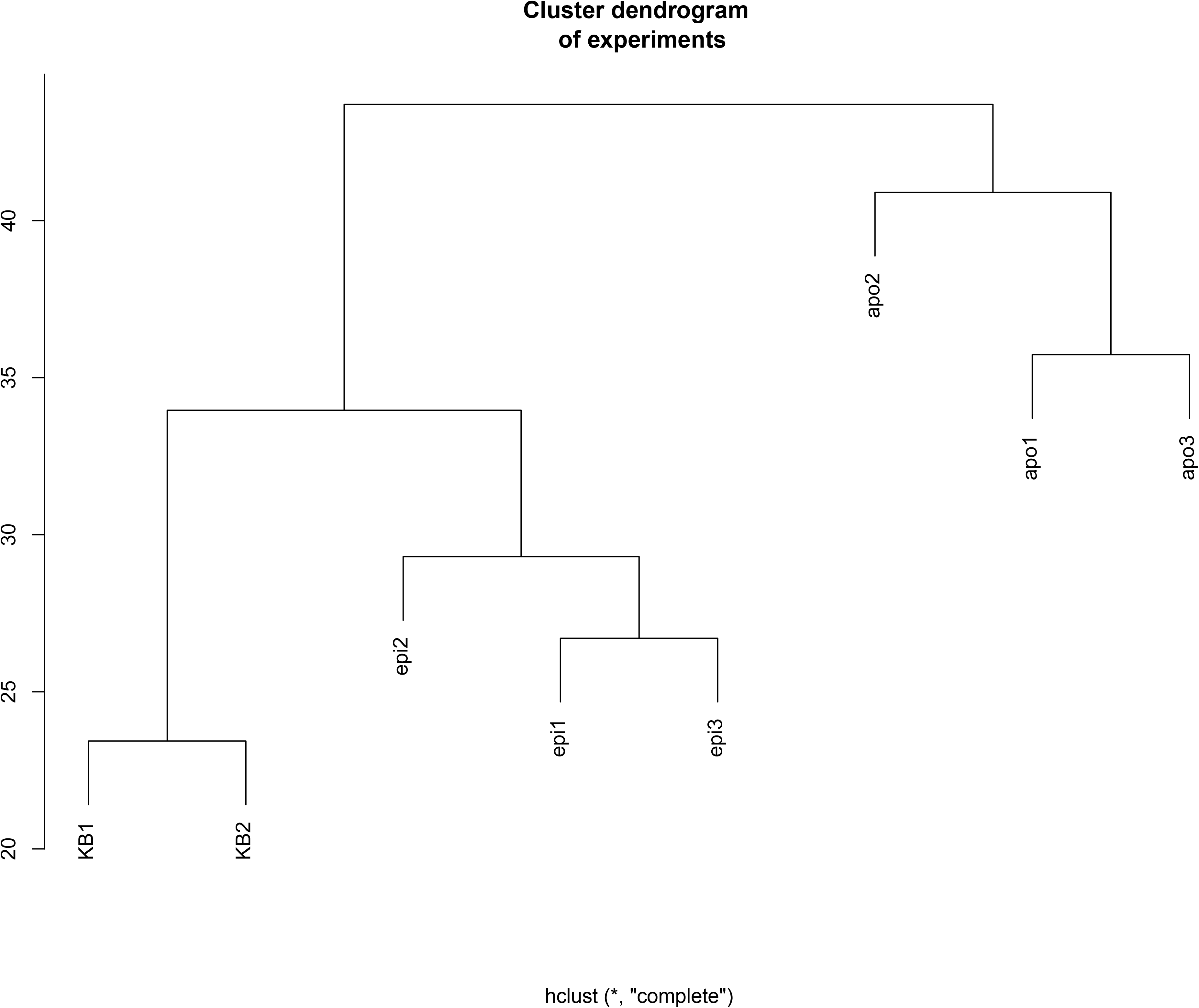
Dendrogram of experiments, generated using *P. syringae* gene fitness scores determined in experimental replicates of the three conditions tested. Rich media King’s B (KB) experiments cluster more closely with the epiphytic experiments (epi) than the apoplastic experiments (apo).

For each experiment, an aliquot of the mutant library was grown to mid-log phase in KB (ca. 5 generations) and a sample of the library was taken immediately before inoculating either the surface or interior of plants (time0). After growth in each condition, cells were recovered from either the surface or interior of the plants and prepared for sequencing. Fitness for each strain was calculated as the log_2_ of the ratio of barcode abundance following growth in or on plants with that barcode abundance obtained initially at time0. Gene fitness is calculated as the weighted average of the individual strain fitness scores (42). Insertions in the majority of genes did not change fitness as measured by relative barcode abundance in the population, and thus the fitness scores for most genes were close to 0.

### Identification of the essential gene set of B728a

Of the 920 genes for which fitness could not be calculated due to a lack of sufficient insertional mutants, only 7 do not contain TA dinucleotide sites and thus are not accessible by mutagenesis with the *mariner* transposon we used. 512 of these genes did contain at least one mapped insertion, but we were unable to calculate fitness scores for them due to the small number of sequencing reads at time0 (Fig. S2), suggesting that they were relatively unfit *in vitro* compared to other mutants, and thus in low relative population size in the library. Based on analysis of the TnSeq data, we predicted 392 genes to be essential for B728a growth on LB, as they contain numerous TA sites but do not contain any mapped insertion strains in our library. The 392 predicted essential genes include many annotated as being involved in translation (including tRNAs), energy generation, and cofactor metabolism (Table S1). We identified homologs in *P. aeruginosa* PAO1 for 363 of the 392 predicted essential B728a genes (Table S2). Of these, 259 are predicted to be essential and 104 are predicted to be nonessential in that strain (44).

### Disruption mutants with fitness defects in rich media

We identified 20 genes that were required for maximal growth in KB media, 9 of which are involved in cofactor metabolism (Table S3). Thirteen of these 20 genes contribute to fitness in or on plants (Fig. 3). This suggests that cofactors such as biotin are lacking in KB media, as well as in the *in planta* habitats.

### Maintaining genetic diversity of the transposon library *in planta*

A major challenge for the use of complex mixtures to study the relative fitness of component strains in any experiment, especially studies done *in planta*, is to ensure that all of the mutants in the mixture are well represented after inoculation so as to avoid bottleneck effects. The strength of saturated transposon mutagenesis methods lies in internal replication: the contribution of each gene is assessed by interrogation of the behavior of multiple independent insertional mutant strains. A loss of diversity at the time of inoculation reduces the statistical power for analysis of a given gene. We aimed to maximize the total number of inoculated bacterial cells to maintain population diversity, while achieving a sufficiently low initial inoculum in or on plants so that substantial, competitive growth of the mixture could be assured. Although we observed slight bottlenecks given the concentration of inoculated cells we used, particularly in apoplastic conditions, these samples provided sufficient reads for most mutants to enable analysis of the fitness contribution of nearly all genes (Table S4); more than 80% and 68% of the unique barcoded mutants were retained in studies of epiphytic and apoplastic growth, respectively. More than 99% of the unique barcoded mutants in the library were retained during *in vitro* experiments.

During the growth of strain B728a on leaf surfaces for 2 days the total number of cells increased approximately 100-fold (27), corresponding to 6 to 7 population doublings. Similarly, during growth in the apoplast for 6 days population size increased about 1000-fold (Fig. S3), indicating at least 10 cell divisions. In theory, in an experiment on leaves in which most strains exhibited 6 generations of growth, mutants completely incapable of growth should exhibit a fitness of about −6 (42). In practice, insertions in very few genes exhibited such an extreme lack of fitness (Fig. 2). However 80 genes in which mutants exhibited fitness scores < −2 (exhibiting only 25% as much growth in the population relative to that of the typical strain in the mutant population) contributed strongly to growth in a given condition (Fig. 3a). Mutants in an additional 69 genes exhibited fitness scores less than −1 but greater than −2 (Fig. 3b) suggesting that these genes contributed somewhat less to fitness (42). We did not normalize fitness scores by the average number of generations in a given experiment, as these values are difficult to estimate and likely vary by plant within an experiment.

**Fig. 2.**
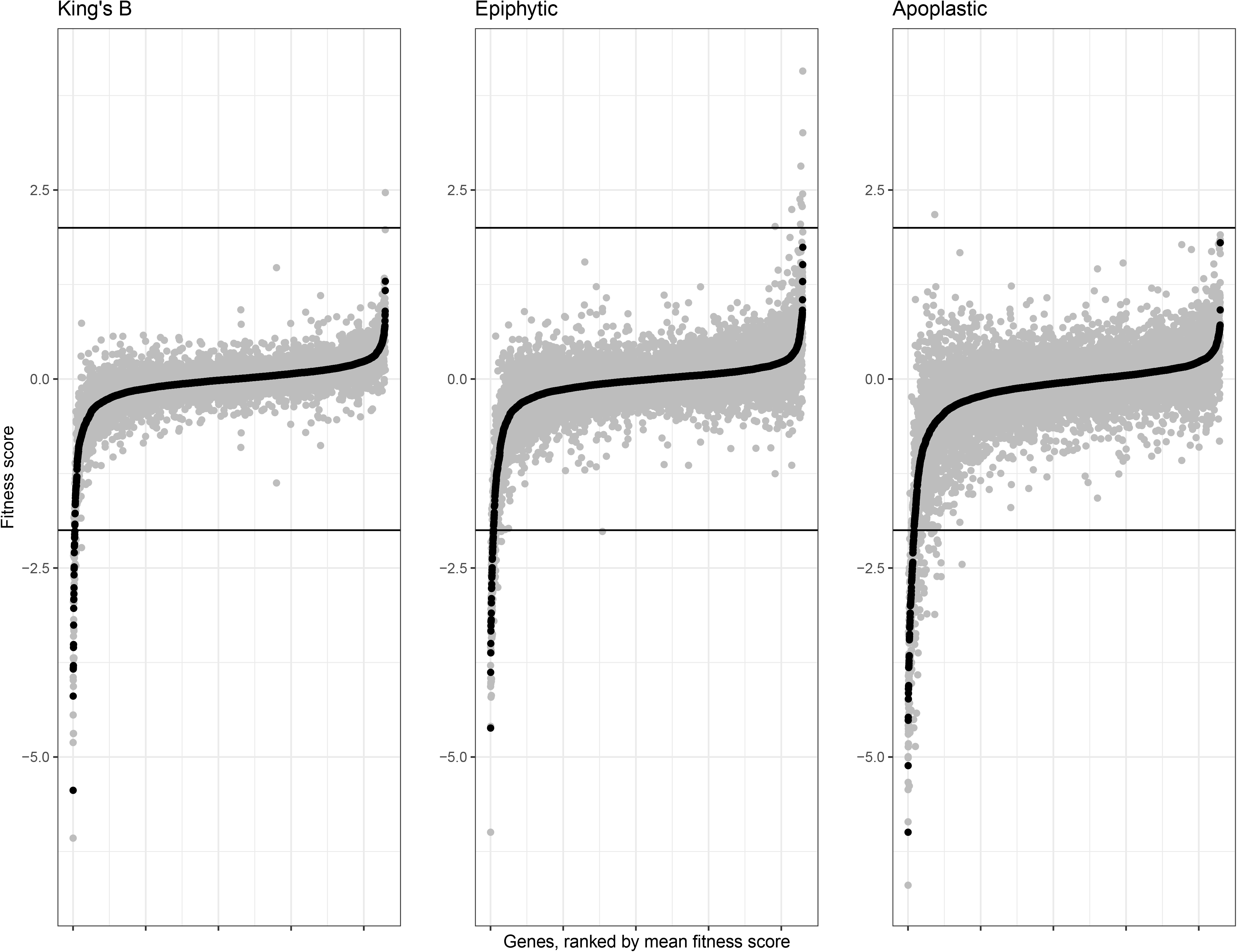
Rank ordered mean gene fitness scores for each condition in which *P. syringae* was grown. Fitness values for independent replicate experiments are shown in grey, while mean fitness scores are plotted in black. Gene fitness scores are calculated as the log_2_ ratio of the barcode counts following growth in a given condition compared to the barcode counts before inoculation. Black lines at fitness values of −2 and 2 are used to indicate strong phenotypes; for example a value of −2 indicates that mutants in that gene were 25% as fit as the typical strain in the mutant library. In each dataset, fitness values < −2 or > 2 are more than 3 standard deviations from the mean (approximately 0).

**Fig. 3.**
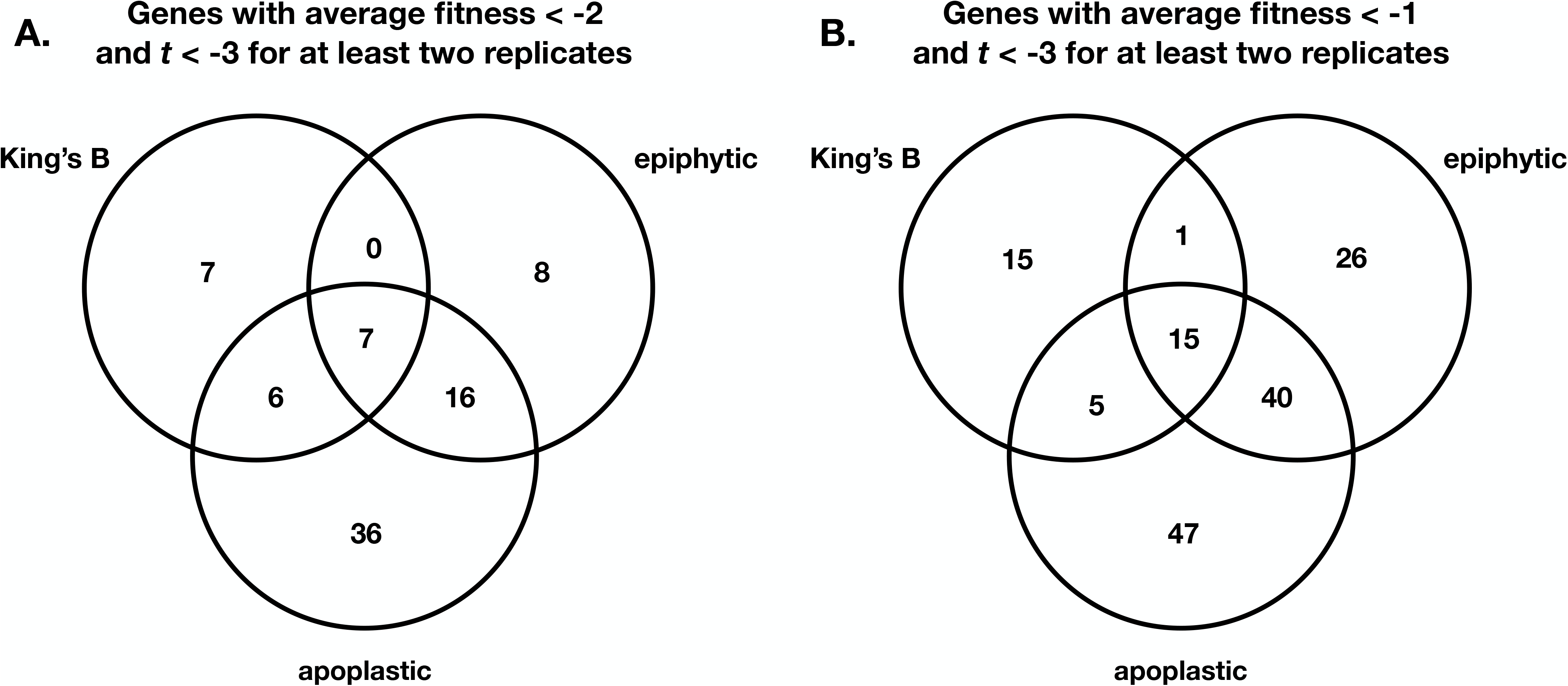
Genes with significant contributions to competitive fitness in the experimental conditions tested. (A) Venn diagram of genes with average fitness values < −2, and *t* < −3 for at least two experimental replicates. (B) Venn diagram of genes with average fitness scores < −1, and *t* < −3 for at least two experimental replicates.

Despite the differences in the number of generations in epiphytic versus apoplastic growth in plants (27), we observed similar ranges in overall fitness scores for individual genes in these two habitats (Fig. 2). Mean fitness scores ranged from −4.6 to +1.7 (epiphytic) and −6.0 to +1.8 (apoplastic). For each gene, we averaged fitness values for the 2 replicate growth experiments in KB and the 3 epiphytic and 3 apoplastic experiments performed. We focused our analysis on genes contributing most strongly to fitness - those having an average fitness < −2 and for which the *t*-score was < −3 in at least two replicate experiments. Since the plant host constitutes a more variable environment than most *in vitro* experiments, and expecting that many genes would not individually make large contributions to fitness, we also examined genes with fitness scores < −1 but with *t* < −3. Special attention was placed on those genes with such scores that are operative in a given metabolic pathway or could be placed in the same functional category. Approximately 50% of all genes exhibiting fitness scores less than − 2 or −1 in either epiphytic or apoplastic habitats were verified in at least two of three replicate experiments (Fig. S4).

### Genes required specifically for colonization of the leaf surface

We identified 31 genes that were highly important for fitness on the leaf surface (Table S5), although only 8 were not also important in the apoplast. Among these 8, genes in the predicted operon *Psyr_2461-2* had a particularly strong epiphytic phenotype, with average fitness scores of −2.1 and −3.2. *Psyr_2461* is a hypothetical protein containing a domain of unknown function (DUF934) and *Psyr_2462* is homologous to the sulfite reductase *cysI* in *P. aeruginosa*. Glutamate synthase (NADPH) subunit genes *gltB* (*Psyr_0411*) and *gltD* (*Psyr_0412*) also contributed strongly to epiphytic growth, having average fitness scores of −2.0 and −1.1. Disruption of the putative phage-related protein Psyr_4512 also strongly reduced epiphytic fitness (average fitness score = −2.1).

### Genes contributing specifically to colonization of the leaf apoplast

Disruption of many genes encoding known virulence factors, including those in the type III secretion system (T3SS) (Fig. S5) and phytotoxin biosynthesis genes greatly reduced the growth of *P. syringae* in the apoplast. Of the 65 genes that were highly important (average fitness < −2) for apoplastic colonization (Table S6), 36 were important in this habitat but not on leaf surfaces. The T3SS genes we observed as essential for successful apoplastic colonization are exclusively involved in the T3SS machinery, as transposon insertions in most individual effector genes generally had no fitness phenotype (Fig. 4a). Of the secreted type III effectors, *hopAB1* had the largest negative average fitness value (−0.55). While the fitness contribution of this gene was less than many others, growth of mutants in this gene was decreased in all three experimental replicates (standard deviation = 0.065, *t* < −3.5 for all). As *t*-values are positively correlated with measures of fitness, the low variance in fitness seen among the 21 insertional mutants for this gene provide confidence in the rather modest fitness estimates for this gene.

**Fig. 4.**
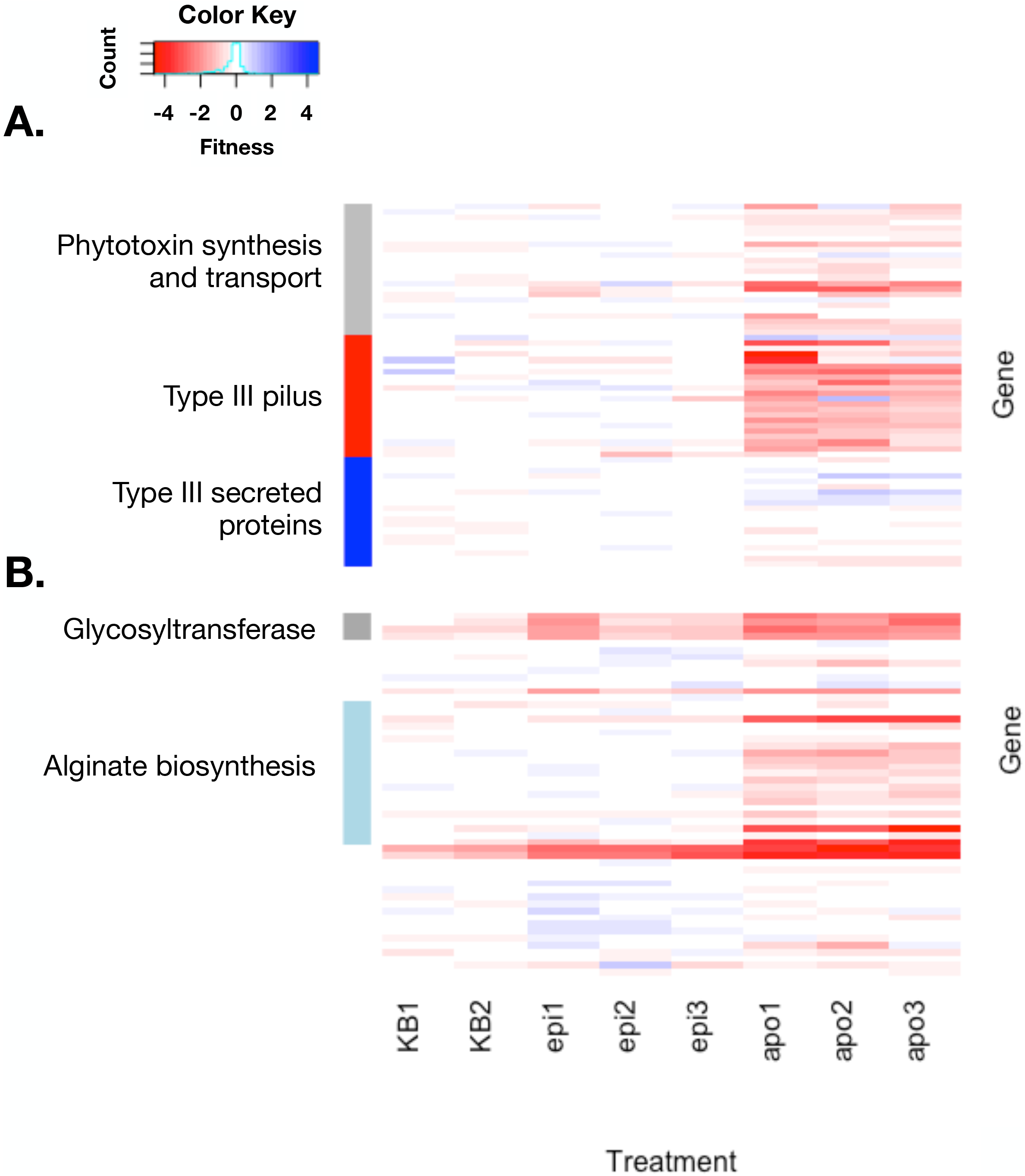
Fitness contributions of genes involved in phytotoxin synthesis and transport and the type III secretion pilus (A) as well as alginate biosynthesis (B) are required for apoplastic colonization. An expanded version of this figure containing gene names and loci can be found in the Supplemental Material.

Production and secretion of the phytotoxin syringomycin was required for competitive fitness in the apoplast. The syringomycin regulator *syrP* and syringomycin efflux transporter *syrD* both had large apoplastic-specific phenotypes when disrupted. In contrast, syringopeptin and syringolin mutants did not have significant apoplastic fitness defects in our experiments. Polysaccharide synthesis and regulation was highly important for competitive fitness in the apoplast. Mutants in alginate regulation (*algU*) and biosynthesis were dramatically less competitive than the typical mutant in the library. Group 1 glycosyltransferase encoding genes (*Psyr_0920* and *wbpYZ*) also contributed substantially to apoplastic growth (Fig. 4b).

The two-component system GacA/GacS was moderately important in the apoplast (average fitness = −0.9 and −1.5), but interestingly, their disruption resulted in increased fitness on the leaf surface (average fitness = 1.3 and 1.7). Conversely, glutathione synthase (*gshB*) was important in KB (average fitness = −1.5) and on the leaf surface (−1.2), but disruption of this gene increased competitive fitness in the apoplast (+1.8). Generally, however, insertional mutations rarely significantly increased fitness in any experiment.

### Genes required for the colonization of both the leaf surface and apoplast

Overall, the categories of “amino acid metabolism and transport”, “polysaccharide synthesis and regulation”, and “nucleotide metabolism and transport” were enriched in genes with average fitness less than −2 in both epiphytic and apoplastic habitats relative to that in rich media (Table S7). We identified 31 genes that were highly important for epiphytic colonization, and 65 genes that contributed to apoplastic growth. Approximately 1/3 of all genes contributing to leaf colonization were also important in the apoplast (Fig. 3).

Genes required for the biosynthesis of several different amino acids were highly important in colonization of both the leaf surface and the apoplastic space. Genes required for biosynthesis of tryptophan, proline, and the shared biosynthetic pathway of isoleucine/leucine/valine were among those with the largest contributions to fitness in both *in planta* conditions tested. Additionally, genes involved in biosynthesis of methionine were important for epiphytic survival, as seen previously (45), and disruption of these genes caused even greater decreases in apoplastic growth suggesting that these resources are in low abundance in these habitats. For example, average fitness scores for *metW* and *metZ* auxotrophs were < −4 in the apoplast, but approximately −0.8 on leaf surfaces. A similar, albeit less dramatic, pattern of proportionally larger requirements for histidine biosynthesis under apoplastic growth was also seen. The production of cofactors such as pantothenate (vitamin B_5_) requiring *panC* was important in both *in planta* conditions but contributed more to the growth on the leaf surface. Genes involved in nucleotide biosynthesis (*purFL*) were also highly important for growth both in and on leaves.

The genes *mdoGH* encoding glucan synthesis were required for optimal growth on both the leaf surface and in the apoplast. Hypothetical proteins encoded by *Psyr_0532*, *Psyr_2461*, and *Psyr_4158* (*eftA*) all made significant contributions to fitness both epiphytically and in the apoplast. Psyr_0532 contains a group 1 glycosyltransferase domain.

### Validation of fitness estimated in disruption mutant mixtures with targeted deletion strains

To determine whether the growth deficiencies of mutant strains in the pooled assays were predictive of that when grown in isolation, we constructed targeted deletion mutants of several genes that contributed differentially to apoplastic fitness of *P. syringae*. Amino acid auxotrophs Δ*trpA* (average apoplastic fitness = −2.7) and Δ*hisD* (average fitness = −3.0) inoculated individually into the apoplast. While Δ*trpA* was almost incapable of growth, the population size of Δ*hisD* was about 10-fold lower than the WT strain 4 days after inoculation. Similarly, while a Δ*hrpL* type III secretion mutant (average fitness = −2.2) achieved an apoplastic population size that was only about 1% that of the WT strain, the population size of a Δ*syrP* deficient in production of syringomycin (average fitness = −2.1) was only slightly lower than that of the WT strain when inoculated separately into plants (Fig. 5a).

**Fig. 5.**
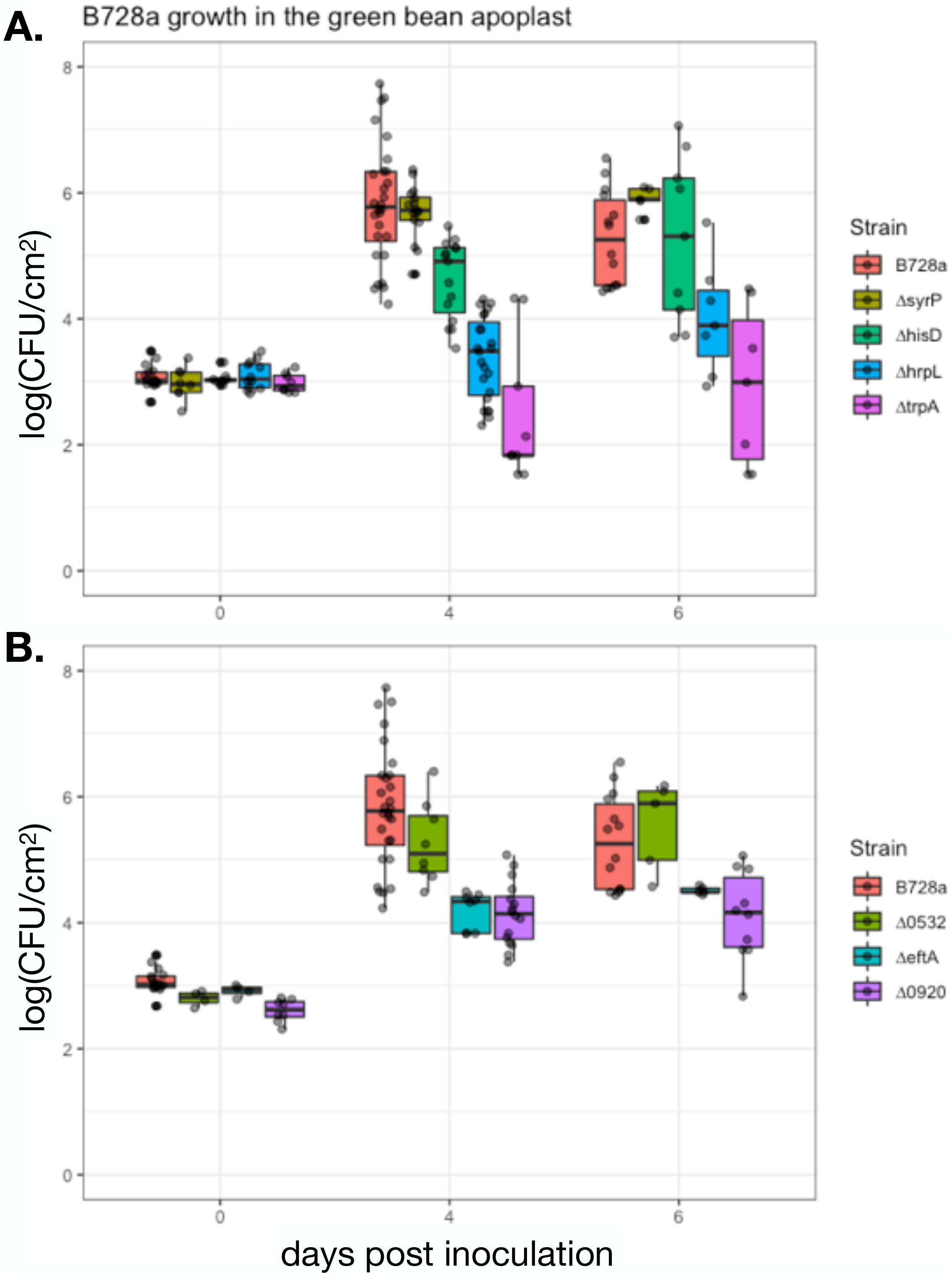
Apoplastic growth of B728a and deletion strains in bean. (A) Growth of the amino acid auxotrophs Δ*trpA* and Δ*hisD*, type III regulatory mutant Δ*hrpL*, and syringomycin mutant Δ*syrP*. At 4 dpi, Δ*hisD*, Δ*hrpL*, and Δ*trpA* are significantly lower than WT B728a. At 6 dpi, Δ*hrpL* and Δ*trpA* are significantly different from WT (Welch Two Sample t-test, p < 0.01). (B) Apoplastic fitness of deletion mutants of glycosyltransferase genes Psyr_0532 and Psyr_0920, and hypothetical protein *eftA*. At both 4 and 6 dpi, the population size of Δ*eftA* and Δ*0920* are significantly lower that WT B728a (Welch Two Sample t-test, p < 0.01).

We also assessed fitness of directed mutants of three poorly understood genes, two of which are predicted group 1 glycosyltransferases, that contributed substantially to competitive apoplastic growth. Deletion mutants of Δ*eftA* (average fitness = −1.4, a hypothetical protein), and Δ*Psyr_0920* (average fitness = −2.4, a group 1 glycosyltransferase) both achieved apoplastic population sizes that were 10-fold lower than that of the WT strain when assessed both 4 and 6 days after inoculation (Fig. 5b). In contrast, the apoplastic population size of Δ*Psyr_0532* (average fitness = −1.6, a hypothetical protein containing a group 1 glycosyltransferase domain), was only slightly less than that of the WT strain (Fig. 5b).

### Fitness contributions of genes do not correlate well with their level of transcriptional expression or inducibility in or on plants

We compared previously published global transcriptional patterns for the genes in strain B728a when grown on leaf surfaces and in the apoplast (27) with that of the fitness values of these genes measured here to determine how predictive gene expression was to the functional role of these genes in growth in an on leaves. Both the absolute levels of gene expression in various *in planta* conditions as well as that of the changes in expression of a given gene *in planta* relative to that in cells grown in a minimal medium were used as predictors. In general, while many genes exhibited substantial elevated or depressed expression on or in plants compared to that in culture media, disruption of these same genes often had little or no impact on the competitive fitness of the mutant strain in this study (Fig. 6). For example, while many amino acid auxotrophs were significantly less fit on the leaf surface and leaf interior, expression of biosynthesis genes for tryptophan, histidine, proline, and methionine was not induced, and instead was repressed, in these habitats compared to *in vitro* conditions. Similarly, while genes involved in biosynthesis of the cofactor pantothenate (shared with biosynthesis of the branched amino acids) were required for competitive fitness, their expression was down-regulated *in planta*. Likewise although the expression of genes encoding several hypothetical proteins were strongly increased *in planta*, suggesting that they may play an important role in growth on plants, their disruption had little effect on the competitive fitness of these mutant strains. Exceptions to this lack of association between gene induction and contribution to fitness are the genes (*syrP* and *syrD*) required for the biosynthesis of syringomycin in the apoplast; these genes were highly up-regulated *in planta* and mutants in this gene cluster also were much less fit. Likewise, *scrB*, involved in sucrose metabolism, is strongly up-regulated specifically in the apoplast and was also specifically required for competitive fitness in that environment. Generally, however, examples of genes in which a concordance between absolute or plant-dependent levels of transcription and their fitness contribution *in planta* were rare.

**Fig. 6.**
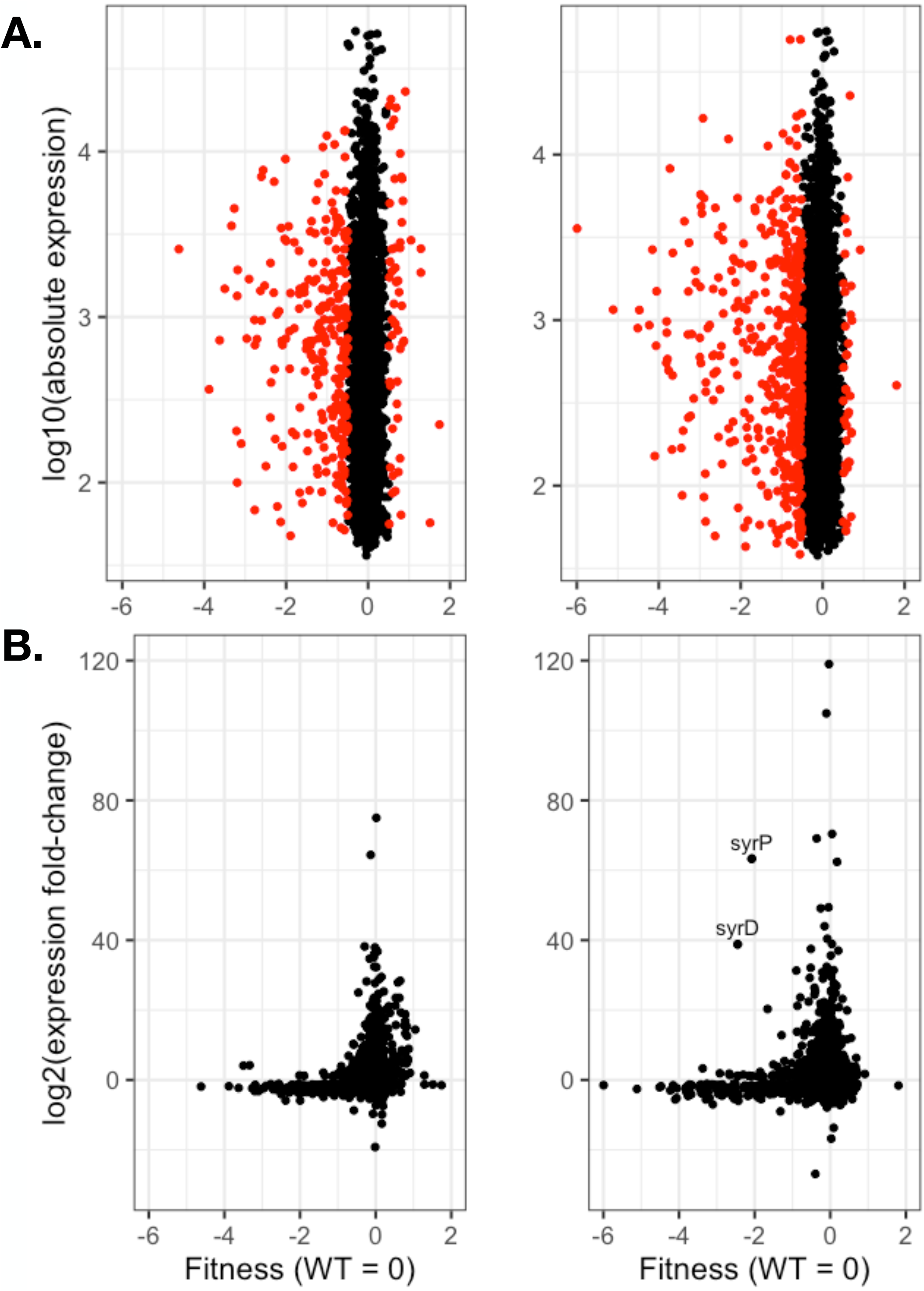
The magnitude of fitness contributions of genes in *P. syringae* do not correlate well with their absolute level of expression (A) or fold-change of these genes *in planta* compared to that in a minimal medium (B). Absolute expression is a measure of fluorescence in microarrays (27). Values of average fitness of mutants either less than - 0.5 or greater than 0.5 are highlighted in red. Fold change in gene expression was calculated as a log_2_ of the ratio of gene expression estimated from microarray fluorescence *in planta* relative to that in a basal medium (27).

## Discussion

Competitive colonization assays are a very sensitive method by which differences in relative fitness can be assessed. In a phenotypically heterogeneous population, changes in relative proportion of a given member provide a direct assessment of relative fitness. A notable exception that would preclude such a process would be one in which there is the production of shared goods (such as siderophores) that can be co-opted by non-producers, as predicted by the Black Queen Hypothesis (46). Random mutagenesis methods such as TnSeq, in which insertional mutant strains are grown in large pooled mixtures, have the advantage of identifying conditionally important genes in a genome-wide manner. By being intrinsically parallel in their structure, the ability to readily distinguish and enumerate each of the individuals in such a mixture by RB-TnSeq provides both a high throughput and highly sensitive means by which relative fitness of the individual strains can be assessed. Furthermore, the creation of multiple independent insertional mutants for each gene provides substantial internal replication, increasing confidence in the fitness phenotype quantified for any given gene. An advantage that RB-TnSeq has over more classical TnSeq is that the association of a given transposon insertion within a gene need be done only once, since a random DNA barcode can then be unambiguously associated with that insertional event thereafter. Such a process then allows the use of the same barcoded transposon library for multiple experiments by simply sequencing and enumerating the DNA barcodes, enabling repeated interrogation of the role of the genes in a species such as *P. syringae* in many different environmental settings. The utility of RB-TnSeq has been demonstrated by its application to a myriad of different bacterial species exposed to hundreds of distinct environmental settings, enabling functions to be assigned to many previously uncharacterized genes (47). Our demonstration of the utility of RB-TnSeq in this study should enable us and others to greatly expand the association of genes in *P. syringae* to the myriad of functions in which it might be expected to participate, in the many chemically and physically different habitats that it colonizes.

*P. syringae* is a robust colonizer of both leaf surfaces and the apoplastic space of the host plant green bean. In these habitats, this strain exhibited sufficient growth to enable RB-TnSeq to quantify the contribution of individual genes that directly contribute to competitive fitness in a heterogeneous population. It would be expected that the ability of such a method to resolve differences in fitness contributions of these genes would increase with the number of generations of growth that the population of mutants would have undergone during an experiment. Given the large number of genes for which some fitness contribution could be measured, we focused our analysis here on those genes having the largest fitness contribution. Furthermore, there is generally higher statistical support for the validity of fitness estimates for those genes contributing more to fitness (Fig. S5), given that they were large and reproducible across replicate experiments. Genes associated with somewhat lower, but consistent, fitness values (Table S8) are likely also biologically significant, and future studies can explore the roles of these genes during *in planta* growth in more depth. In the current study the high internal replication intrinsic to the barcoded transposon library, and the use of several replicate experiments for each *in planta* condition has provided a compelling list of broadly important genes for further analysis.

Transposon-based approaches have been useful in identifying essential bacterial genes in other taxa, although our knowledge of essential genes in *Pseudomonas* species is limited. Analysis of a 100,000 strain insertional library in *P. aeruginosa* identified 336 of the 5,606 genes to be essential (44). Recent work has identified 473 genes as likely to be essential in *P. simiae*, 430 in *P. stutzeri*, and 325 to 442 in *P. fluorescens* (depending on the strain) (47). We identified 392 genes to be likely essential for B728a growth in LB, comprised of functional categories generally seen to be essential in diverse bacteria. Given that we could calculate fitness contributions for 84% of the protein-coding genes in the environmental conditions tested here, the proportion of genes found to be essential in *P. syringae* appears similar to that *E. coli* and *P. stutzeri* (42).

The identification of genes involved in anabolic processes such as cofactor production and amino acid biosynthesis that contribute to the fitness of *P. syringae* in a given habitat provides some insight into the availability of such resources in that setting. This logic of anabolic mutants as reporters of habitat resources provides insight into the resources on the surface and in the intercellular spaces of plants. The finding of fitness defects for many amino acid biosynthetic genes is a clear example of this concept. The much lower fitness of auxotrophs for several different amino acids suggests an acute limitation of these essential metabolites both on the leaf surface and in the apoplast. Genes within the biosynthetic pathway of tryptophan had the largest effect on fitness when disrupted in our study, both on the leaf surface and in the apoplast. The biosynthesis of tryptophan is more energetically costly than other amino acids in *E. coli* (48). Additionally, tryptophan is utilized for downstream biosynthetic pathways in B728a such the synthesis of auxin, a plant hormone shown to contribute to epiphytic fitness of some bacteria (8). Similar to our observations, biosynthetic genes for tryptophan (and its precursor anthranilate) were identified as important for fitness in a TnSeq study examining *Pantoea stewartii* colonization of maize xylem (49). A TnSeq screen in *Dickeya dadantii* in rotting plant tissue also noted a significant decrease in competitive fitness *in planta* for leucine, cysteine, and lysine auxotrophs that could be negated through the external addition of amino acids (50). Since most amino acids are apparently present at relatively low concentrations in the bean apoplast (9), it could be expected that many auxotrophs are incapable of growth without the ability to synthesize these essential non-substitutable metabolites, a model supported by the observations of this study. Indeed, bacterial biosensors are often used *in situ* as an alternative to direct metabolite measurement to detect diverse environmental conditions, such as sugar availability (4).

Unlike certain anabolic genes, those involved in central metabolism typically had more subtle phenotypes when disrupted. This is likely due to the presence of diverse and substitutable carbon and nitrogen sources such as sugars and organic acids in and on plants (9). It was noteworthy that the fitness of mutants in which sucrose 6-phophate hydrolase encoded by *scrB* was disrupted was lower in the apoplast (fitness score −1.7). Such an observation is consistent with sucrose being the most abundant sugar in intercellular spaces (9). On the other hand, the genes involved in the metabolism of compounds of lesser abundance that that are not essential would be expected to contribute somewhat incrementally to the fitness of *P. syringae* in or on leaves. While carbon availability appears to limit the growth of bacteria such as *P. syringae* on leaves (6) and might also limit the growth of this species in the apoplast, it might be expected that these various compounds represent substitutable resources (9), and that any given compound would contribute relatively little to the overall growth of such a strain if many were present in similar concentrations. In support of this conjecture, while small fitness defects were observed for several mutants harboring disruption of genes essential for catabolism of nutritive compounds, the magnitude of these fitness defects was usually low.

Many genes involved in polysaccharide synthesis and regulation were highly important in leaf colonization. The lipopolysaccharide found in the outer membrane of Gram-negative bacteria is known to induce the innate immune response of plant and animal hosts, yet it is required for bacterial stress tolerance in diverse environments (51). Many of the hypothetical proteins having strong plant phenotypes when disrupted here contain glycosyltransferase domains. We hypothesize that these hypothetical proteins contribute to the biosynthesis of O-antigen that decorates LPS, and thus might be involved in camouflaging the cells so as to not be perceived by plant surveillance systems. O-antigen is an essential virulence factor for *P. aeruginosa* in its colonization of animal tissues (52). While O-antigen is dispensable for growth in culture, it has been recently shown to delay the host immune response during *Xylella fastidiosa* colonization of grape xylem (53). Alternatively, glycosyltransferase activity may contribute to flagellar modifications in order to avoid plant recognition (54, 55). However, this is unlikely to be the major role of these selected genes in *P. syringae* since known flagellar glycosyltransferases in *P. syringae* (*fgt1* and *fgt2*) located adjacent to flagellar biosynthesis genes did not measurably contribute to competitive fitness *in planta* in this study. Nonetheless, we show that 8 genes containing glycosyltransferase domains made large individual contributions to host colonization, suggesting that there may be other important targets for such modification.

Biosynthesis of the exopolysaccharide alginate contributed strongly to growth in the apoplast but not on leaf surfaces. While alginate had been shown to contribute to epiphytic fitness and thus to subsequent disease severity of *P. syringae* (56, 57), the apoplastic colonization of mutants was not distinguished from epiphytic growth in those studies. The apoplast is thought to be a water-limiting environment for endophytic pathogens (10, 12, 27). Furthermore, the transcriptional activation of the key alginate biosynthetic enzyme *algD* is induced by high osmolarity (58) and thus alginate biosynthesis would be expected to contribute to fitness in the apoplast, as observed here. In *P. putida*, alginate production is required for biofilm-mediated survival under desiccating conditions (59). While we did not see a significant role of alginate on the leaf surface, its biosynthesis is a clear virulence factor in the apoplast.

Our screen highlighted the fitness role of many known virulence factors including the type III secretion system and the phytotoxin syringomycin. Individual secreted effector proteins did not generally contribute measurably to apoplastic colonization, while mutations in type III pilus genes significantly decreased fitness. This supports existing dogma, whereby type III effector proteins are individually dispensable and collectively essential (60). HopAB1, a secreted type III effector which we found to have the largest contribution to apoplast fitness among all secreted effectors, has been shown to have a measurable contribution to B728a growth in the bean apoplast (20). While phytotoxin production in strain B728a has been shown previously to induce symptom formation, there has not been compelling data showing a contribution to bacterial growth in plants (61). It is interesting that the genes involved in syringomycin and syringopeptin have strong negative fitness values in our study, suggesting that mutants in these pathways are impaired in growth relative to wild type. Our results support the model that *P. syringae* strains such as B728a with relatively fewer type III effectors have an increased reliance on phytotoxin production for growth in the apoplast (23, 62). The biosynthetic gene cluster for syringomycin also was distinctive in that it was among the few genes that are both up-regulated *in planta* (15) and contribute to fitness. On the other hand, genes for syringolin biosynthesis while up-regulated in the apoplast (27), did not contribute to apoplastic fitness in our study. Syringolin contributes to virulence through host proteasome inhibition, which has been shown to counteract stomatal innate immunity (63). Therefore, the role of syringolin is likely limited to the transition from epiphytic to apoplastic growth, a process that was not tested here. Syringopeptin, which is also up-regulated in the apoplast (27), contributed to apoplastic fitness to a much lesser extent than syringomycin. Syringomycin and syringopeptin have the same mechanisms of action, creating membrane pores and causing ion leakage (64). It is unclear why B728a produces two seemingly redundant phytotoxins, although it has been proposed that their differential antimicrobial activities contribute to epiphytic survival (65). Both syringomycin and syringopeptin contribute to virulence on cherry (64) and lysis of tobacco protoplasts (65). Since we observed a much larger contribution to bacterial fitness in the bean apoplast from syringomycin than syringopeptin, and it is tempting to speculate that the functions of these phytotoxins in virulence may be somewhat host specific.

Despite the dogma that gene expression is fine-tuned to the metabolic demands of a cell, recent studies of gene expression have shown such regulation to be suboptimal for many bacterial species (33). Despite classic examples of biosynthetic pathways in *E. coli* having adaptive regulation, many genes in diverse bacteria show little correlation between when they are important for fitness and when they are most highly expressed (33). For example, constitutive expression and regulation by growth rate are common indirect gene regulation strategies that occur for genes with diverse functions and yet are often suboptimal in the laboratory and presumably also in natural environments (33). In *P. aeruginosa* wound infections, gene expression was also not well correlated with gene contributions to fitness (35). A proposed explanation for such incongruence was that given that *P. aeruginosa* is considered an opportunistic pathogen that might not have evolved primarily in association with mammalian tissues its patterns of gene expression might have optimized fitness in very different settings (35). Moreover, in persistent, long-lasting infections such as the cystic fibrosis lung, adaptive changes in global patterns of gene expression in *P. aeruginosa* have been observed over time (66), reflecting adaptation to this new habitat.

While *P. syringae* is a model plant pathogen, it is also commonly observed in many other environmental settings (19). The conditions that the cell would experience on the surface of the plant are likely to be quite different from those in the apoplast (3, 13). Here, we see no correlation between gene expression (either absolute or relative change) and contribution to fitness in the host. While the timing of sampling of RNA from the apoplast for this comparative study was somewhat earlier in the infection process (Yu *et al.* (27) sampled bacterial cells 2 days post inoculation, while we sampled after 6 days), we would not have expected temporal changes in gene expression to overwhelm any context-dependent patterns of gene expression. It was surprising that genes that were highly expressed and/or highly induced in cells in or on leaves did not make large contributions to the fitness of the strain. Likewise, many genes that were either weakly expressed or un-induced on or in plants proved particularly important for fitness in these habitats. This lack of congruence can be explained by the fact that many genes are involved in catabolic processes wherein individual pathways would be expected to contribute only incrementally to the success of a strain. Genes for anabolic pathways, on the other hand, might prove essential irrespective of how highly expressed they are. There remain many genes for which the lack of linkage between expression and contribution to fitness remain unexplained. It is evident that directly measuring the contribution of a gene to fitness in different environments is a necessary complement to global transcriptional profiling to understanding the function and behavior of a cell in a given setting.

Although *P. syringae* is a ubiquitous species, it is most commonly studied in its agriculturally relevant, disease susceptible plant hosts. Random mutagenesis studies typical observe that a majority of genes in the genome are dispensable, as seen in the relatively small number of essential genes across diverse bacteria (67). This is likely due to many genes contributing to bacterial fitness in untested habitats outside of the laboratory (47). Although previous transposon screens in *P. syringae* have provided information on traits required for epiphytic fitness and virulence, these have either uncovered only those genes with large effects on behavior, or which could be readily performed *in vitro* (68, 69). RB-TnSeq greatly expands our ability to interrogate the ecological determinants of such a cosmopolitan bacterium. Testing *P. syringae* and other bacterial species in a range of conditions, including those of ecological relevance such as on and in additional host and non-host plants, will enable the designation of functions for hypothetical or otherwise uncategorized proteins. Comparisons of these fitness assessments with specific *in vitro* experiments will enable the dissection of how individual genes contribute to a given process and to fitness on a eukaryotic host, a complex habitat with many distinct abiotic and biotic stressors. In such an approach, Cole *et al.* used this method to examine specific nutrient requirements for *P. simiae* colonization of *Arabidopsis* roots (43). Many of the genes found to contribute to fitness had only small effects *in planta*. Expansion of these screens through additional generations of growth will increase the accumulated fitness defects, as seen in a recent study that sequentially passaged a *Caulobacter crescentus* transposon library to identify genes affecting attachment (70). Barcoded transposon libraries were originally developed as a highly scalable tool to identify gene function in diverse *in vitro* conditions such as different growth conditions or abiotic stresses. Here we show that these same libraries can be used to better understand conditionally important genes that contribute to growth on the leaf surface and during colonization of the apoplast, expanding our understanding of the ecological fitness requirements on a genome-wide scale.

## Materials and Methods

### Bacterial strains and growth media

*P. syringae* pv. *syringae* B728a was originally isolated from a bean leaf (*Phaseolus vulgaris*) in Wisconsin (17). The complete genome for B728a is available on NCBI GenBank as accession CP000075.1 (71). B728a and derivative mutant strains were grown on King’s B agar or in broth (72), at 28°C. *E. coli* strains S17-1, TOP10, and XL1-Blue were grown on LB agar or in LB broth at 37°C. When appropriate, the following antibiotics were used at the indicated concentrations: 100 μg/ml rifampicin, 50 μg/ml kanamycin, 15 μg/ml tetracycline, 40 μg/ml nitrofurantoin, and 21.6 μg/ml natamycin.

### Construction of bar-coded transposon library

The bar-coded transposon insertion library was constructed by transposon mutagenesis using a bar-coded *mariner* transposon library, followed by TnSeq mapping and barcode association as previously described (42). The *E. coli* WM3064 donor library containing the barcoded *mariner* plasmid, pKMW3, was recovered from a glycerol stock in LB kanamycin containing 300 μM diaminopimelic acid (DAP) and conjugated into B728a overnight on LB plates containing DAP. The conjugation mixture was resuspended and spread on LB kanamycin plates for selecting mutants. Over 220,000 kanamycin resistant B728a colonies were pooled for the library. All colonies were resuspended in 250 ml LB kanamycin and diluted to a starting OD_600_ 0.2 for outgrowth at 28°C with shaking to OD_600_ 1.0. Finally, 250 μl 80% glycerol was added to 1 ml aliquots and frozen at −80°C.

### Plant growth conditions

Common bean (*P. vulgaris* var. Blue Lake Bush) seeds (5 - 7 per 10 cm diameter pot) were planted in Super Soil and grown in a greenhouse for two weeks before inoculation. Leaves were kept dry to minimize epiphytic contamination.

### Library recovery and growth in KB

For each inoculation, a 1.25 ml glycerol stock containing the transposon library was inoculated from −80°C into 25 ml fresh KB with 100 μg/ml kanamycin and grown for approximately 7 hours at 28°C with shaking until the culture reached mid-log phase, OD_600_ 0.5 - 0.7. Time0 samples were collected at this point during recovery; 1 ml aliquots were pelleted by centrifugation and the pellets were frozen until DNA purification. Cells were then washed twice in 10 mM KPO_4_ prior to plant inoculation.

To assay library growth in KB, 50 μl log phase cell culture (OD_600_ 0.5) was inoculated into 950 μl KB with kanamycin in a 24-well plate. The plate was incubated at 28°C with shaking for 15 hours. Cells were collected by centrifugation, and frozen prior to DNA purification.

### Plant inoculations of the transposon library

For epiphytic inoculations, cells were resuspended to a concentration of 2×10^6^ CFU/ml 10 mM KPO_4_ (OD_600_ = 0.001, by dilution from OD_600_ = 0.1), and sprayed onto the leaf surface until runoff. 100 pots were inoculated for a given experiment. Plants were then placed in a high humidity chamber for two days.

For apoplastic inoculations, cells were resuspended to a concentration of 2×10^5^ CFU/ml 1 mM KPO_4_. The soil was covered with cotton to hold the soil in place, and the pots were inverted in ~1.5 L inoculum in a glass bell jar. A vacuum was applied for 1.25 minutes and then removed rapidly to force the inoculum into the apoplast. Ca. 100 pots were inoculated for a given replicate experiment. Plants were allowed to dry overnight and then moved to the greenhouse for six days.

### Library isolation from the leaf surface

Leaves were collected and placed in a water-filled glass dish placed in a sonication water bath to remove cells. The resulting leaf wash was filtered through a 6 μm filter (whatman #3), and then cells were collected on 0.2 μm filters. Cells were removed from the filters by vortexing in 30 ml total 10 mM KPO_4_, and centrifuged at 17,000 x g for 1 minute to pellet. Cell pellet aliquots were frozen prior to DNA purification.

### Library isolation from the apoplast

Leaves were chopped in a blender and placed in a water-filled glass dish placed in a sonication water bath to remove cells. The resulting slurry was filtered through a coffee filter to minimize plant debris. 10% of the ~5-10 L buffer was taken for additional filtration steps. This sample was filtered through several whatman filters (20 μm, 10 μm, and 6 μm), and then concentrated by centrifugation at 4696 x g for 10 minutes. The pellet was resuspended in water, and aliquots of cell pellets were frozen prior to DNA purification.

### DNA isolation and library preparation

DNA from frozen pellets was isolated using the Qiagen DNeasy Blood & Tissue Kit according to manufacturer’s instructions. Cell lysis was done at 50°C for 10 minutes as per optional instructions. For apoplastic samples with excess plant material, lysed cells were centrifuged at 1,500 x g for 5 minutes before loading the supernatant onto purification columns. Purified genomic DNA was measured on a nanodrop and 200 ng of total DNA was used as a template for DNA barcode amplification and adapter ligation as established previously (42). For each time0 and plant experimental sample, two separately purified DNA samples were sequenced as technical replicates.

### Sequencing and fitness data generation

Barcode sequencing, mapping, and analysis to calculate the relative abundance of barcodes was done using the RB-TnSeq methodology and computation pipeline developed by Wetmore *et al.* (42); code available at bitbucket.org/berkeleylab/feba/. TnSeq was used to map the insertion sites and associate the DNA barcodes to these insertions. Based on the TnSeq data, standard computational methods (47) were used to predict which genes are likely essential for viability in LB. For these data, the minimum gene length to call a gene essential was 325 bp. For each experiment, fitness values for each gene are calculated as a log_2_ ratio of relative barcode abundance following library growth in a given condition divided by relative abundance in the time0 sample. Fitness values are normalized across the genome so the typical gene has a fitness value of 0. All experiments passed previously described quality control metrics (42). Experimental fitness values are publically available at fit.genomics.lbl.gov.

### Comparison of *P. aeruginosa* predicted essential genes to genes lacking fitness data

We used the Integrated Microbial Genomes (IMG) database (73) to identify homologs for B728a genes in *P. aeruginosa* PAO1 using the genome-gene best homologs function. Turner *et al.* predicted 336 essential genes in PAO1 using a Monte Carlo statistical analysis (44). A comparison of B728a genes predicted to be essential (N = 392) with their PAO1 homologs identified three categories: predicted essential and nonessential PAO1 genes, as well as B728a genes with no PAO1 homologs identified.

### Genomic fitness data analysis

A dendrogram of experiments was generated from the matrix of fitness values using the hclust function in R (74) with the default clustering method “Euclidean”. To better classify genes based on their genomic annotation, we assigned gene names, gene product descriptions, and broad functional categories based on the previously annotated genomic metadata (27). For each gene, fitness values for experimental replicates were averaged to calculate an average gene fitness score for each treatment. We focused our analysis on genes with average fitness < −2 and *t* < −3 in at least two experimental replicates. However, we also considered genes for analysis with average fitness < −1 and *t* < −3 in at least two experimental replicates. The *t*-score is a test statistic used to assess the statistical significance of the gene fitness scores (42). For each functional category, we used a hypergeometric test (phyper function in R) to examine category enrichment, using average fitness < −2.

### Construction of targeted deletion mutants

Deletion strains were constructed using an overlap extension PCR protocol as describe previously (75). Briefly, 1kb DNA fragments upstream and downstream the genes of interest were amplified along with a kanamycin resistance cassette from pKD13 (76). These three fragments were joined by overlap extension PCR. The resulting fragment was blunt-end ligated into the SmaI site of pT*sacB* (77), and transformed into the *E. coli* subcloning strains TOP10 or XL1-Blue, and then the *E. coli* conjugation donor strain S17-1. This suicide plasmid was conjugated into B728a on KB overnight, and then selected for 3 days on KB containing kanamycin and nitrofurantoin (*E. coli* counter selection). Putative double-crossover colonies that were kanamycin resistant and tetracycline sensitive were selected for screening using external primers and further confirmed by PCR and Sanger sequencing.

### Bacterial apoplastic growth measurements

Strains were grown overnight on KB, washed in 10 mM KPO_4_, and standardized to 2×10^5^ CFU/ml in 1 mM KPO_4_. Cells were inoculated into leaves of two-week old plants using a blunt syringe. Leaf samples were taken using a 5 mm-diameter cork borer into tubes containing 200 μl 10 mM KPO_4_ and two 3 mm glass beads, and ground for 30 seconds at 2400 rpm in a Mini-Beadbeater-96 (Biospec Products) before dilution plating on KB with rifampicin and natamycin (an anti-fungal).

## Acknowledgements

We thank Morgan Price for assistance with RB-TnSeq sequence analysis, and Jayashree Ray for mapping the insertions sites of the B728a transposon library. We thank Russell Scott for the B728a Δ*hrpL* strain. We thank Dana King, Caitlin Ongsarte, and Jennifer Lam for assistance with plant inoculations. Funding for Tyler Helmann was partially provided by the Arnon Graduate Fellowship and the William Carroll Smith Fellowship. This work used the Vincent J. Coates Genomics Sequencing Laboratory at UC Berkeley, supported by NIH S10 OD018174 Instrumentation Grant.

