## Supplemental Material for "Genome-wide identification of *Pseudomonas syringae* genes required for competitive fitness during colonization of the leaf surface and apoplast"

**Supplementary data and figures**

Strains and plasmids

| Strains | | |
| --- | --- | --- |
| Organism | Description | Source |
| *E. coli* TOP10 | For general cloning | Invitrogen |
| *E. coli* XL1-Blue | For general cloning | QB3 Macrolab |
| *E. coli* S17-1 | Conjugation donor strain | (1) |
| *E. coli* WM3064 | Strain APA752; barcoded *mariner* transposon vector (Kan^R^) in *E. coli* conjugation strain | (2) |
| *P. syringae* B728a | Wild type strain (Rif^R^) | (3) |
| *P. syringae* B728a | Whole genome barcoded *mariner* transposon library (Rif^R^ Kan^R^) | This work |
| *P. syringae* B728a | ∆*hrpL* (Rif^R^) | (4) |
| *P. syringae* B728a | ∆*trpA* (Rif^R^ Kan^R^) | This work |
| *P. syringae* B728a | ∆*hisD* (Rif^R^ Kan^R^) | This work |
| *P. syringae* B728a | ∆*syrP* (Rif^R^ Kan^R^) | This work |
| *P. syringae* B728a | ∆*eftA* (Rif^R^ Kan^R^) | This work |
| *P. syringae* B728a | ∆*Psyr_0532* (Rif^R^ Kan^R^) | This work |
| *P. syringae* B728a | ∆*Psyr_0920* (Rif^R^ Kan^R^) | This work |

| Plasmids | | | |
| --- | --- | --- | --- |
| Plasmid name | Description | Antibiotic | Reference |
| pKD13 | Source of kanamycin resistance | Kan | (5) |
| pT*sacB* | Suicide plasmid to introduce DNA into *P. syringae* | Tet | (6) |
| pT:0033-kan | To delete *trpA*, contains *Psyr_0033* flanking regions bordering kan^R^ cassette, inserted into SmaI site of pT*sacB* | Tet Kan | This work |
| pT:4133-kan | To delete *hisD*, contains *Psyr_4133* flanking regions bordering kan^R^ cassette, inserted into SmaI site of pT*sacB* | Tet Kan | This work |
| pT:2612-kan | To delete *syrP*, contains *Psyr_2612* flanking regions bordering kan^R^ cassette, inserted into SmaI site of pT*sacB* | Tet Kan | This work |
| pT:4158-kan | To delete *eftA*, contains *Psyr_4158* flanking regions bordering kan^R^ cassette, inserted into SmaI site of pT*sacB* | Tet Kan | This work |
| pT:0532-kan | To delete *Psyr_0532*, contains *Psyr_0532* flanking regions bordering kan^R^ cassette, inserted into SmaI site of pT*sacB* | Tet Kan | This work |
| pT:0920-kan | To delete *Psyr_0920*, contains *Psyr_0920* flanking regions bordering kan^R^ cassette, inserted into SmaI site of pT*sacB* | Tet Kan | This work |

Primers

Bold sequence complements FRT-Kan sequence for splicing by overlap extension protocol. The primers used for TnSeq mapping and BarSeq amplification are described in (2).

| Name | Sequence |
| --- | --- |
| FRT-KanF | GTGTAGGCTGGAGCTGCTTC |
| FRT-KanR | ATTCCGGGGATCCGTCGACC |
| 0033 up F | TGATCGGCTGCCCTTATGTG |
| FRT 0033 up R | **GAAGCAGCTCCAGCCTACAC**AGCAAGACACAAGGGGTTCA |
| FRT 0033 dwn F | **GGTCGACGGATCCCCGGAAT**TTTGCATGTCTTTGTCGCCG |
| 0033 dwn R | TGGTGTTAGACCTCAACCGC |
| 4133 up F | CTTCCAGCAACGCCTGATGT |
| FRT 4133 up R | **GAAGCAGCTCCAGCCTACAC**TGAGCAAATTCTGGAGCCCT |
| FRT 4133 dwn F | **GGTCGACGGATCCCCGGAAT**GGGCCTCAATAATTGGCGGA |
| 4133 dwn R | TGATCGACCGCATCTACCAC |
| 4133 up F | CTTCCAGCAACGCCTGATGT |
| 2612 up F | CACCCCAGATTTCCCAGACC |
| FRT 2612 up R | **GAAGCAGCTCCAGCCTACAC**GACCTCAGCCCTTCACATCC |
| FRT 2612 dwn F | **GGTCGACGGATCCCCGGAAT**TCCCGTTATCAAGCCAGGAC |
| 2612 dwn R | TGGAGAATCCGAAATCCGCC |
| 4158 up F | CAGGACTCGGAGATCATGCC |
| FRT 4158 up R | **GAAGCAGCTCCAGCCTACAC**CGCCTCATGGAGTACAGTGG |
| FRT 4158 dwn F | **GGTCGACGGATCCCCGGAAT**GGTGCAAAGAGCAGAATCGG |
| 4158 dwn R | GTATCGACTCGCGGGAAACT |
| 0532 up F | ACCTCGTCTCTGGCTGTTTC |
| FRT 0532 up R | **GAAGCAGCTCCAGCCTACAC**CAGTACTGCGCCTGCTGAAT |
| FRT 0532 dwn F | **GGTCGACGGATCCCCGGAAT**ATGATGGTATTCAGCGAAAACAG |
| 0532 dwn R | TAATCCCGGCCACGACAAAG |
| 0920 up F | GTACGCTGGAAGAATCGGGT |
| FRT 0920 up R | **GAAGCAGCTCCAGCCTACAC**TTTCCTTGCGCTCAAAAGCC |
| FRT 0920 dwn F | **GGTCGACGGATCCCCGGAAT**ACAGCCGATTTGAACCTGGG |
| 0920 dwn R | ACTTCCATGCCAGAAGGTGG |

Fig. 4 Expanded. Fitness contributions of genes involved in phytotoxin synthesis and transport and the type III secretion pilus (A) as well as alginate biosynthesis (B) are required for apoplastic colonization.

A.

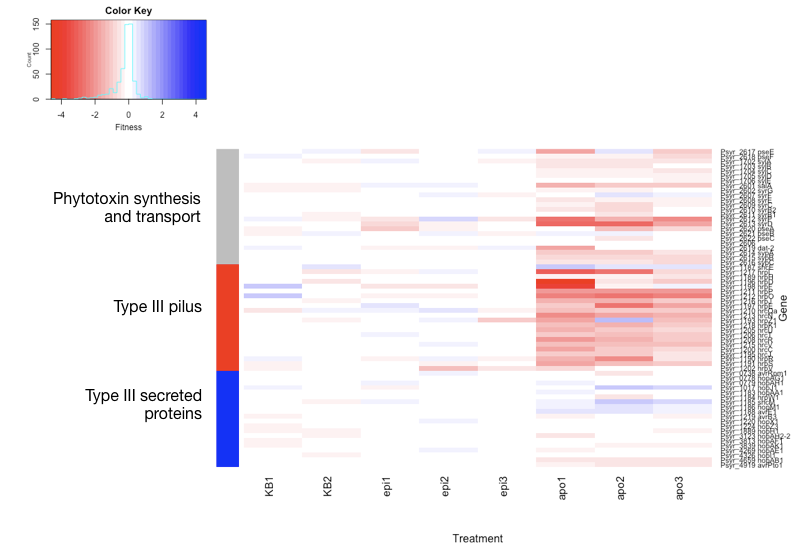

B.

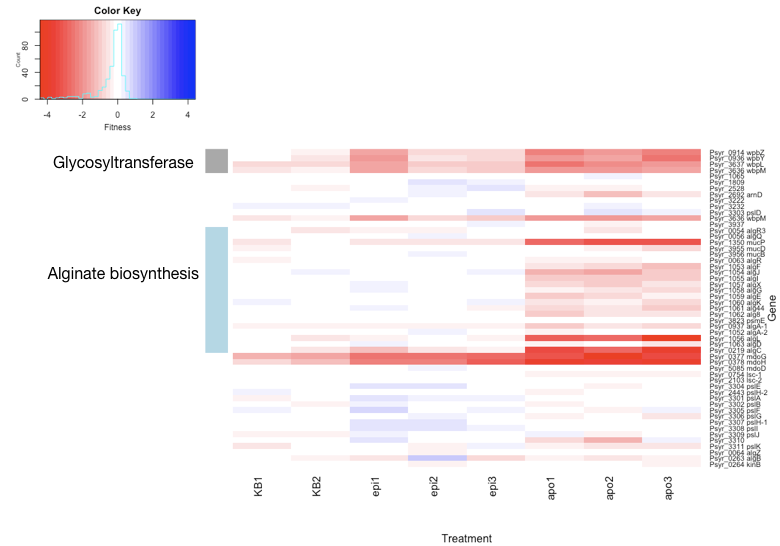

Fig. S1. The B728a *mariner* transposon library contains 281,417 mapped insertions, 169,826 of which lie within the central 10 – 90 % of a gene. Only these central insertions are used to calculate fitness. (A) The number of usable insertions for each gene is correlated with the number of TA dinucleotide sites within each coding region. The Pearson correlation coefficient calculated for all genes having at least 1 usable insertion strain: r = 0.72. (B) 4,296 genes contained at least one usable insertion. Genes for which their contribution to fitness could be calculated are represented by a median of 24 insertion strains per gene. 38 genes with > 200 insertions each (range = 203 to 676) are plotted at 200 for clarity.

A.

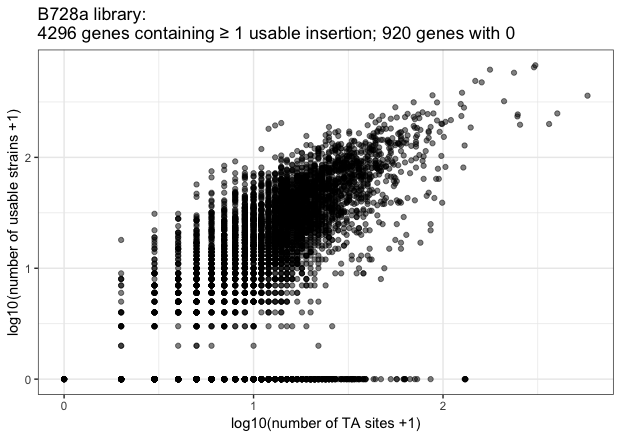

B.

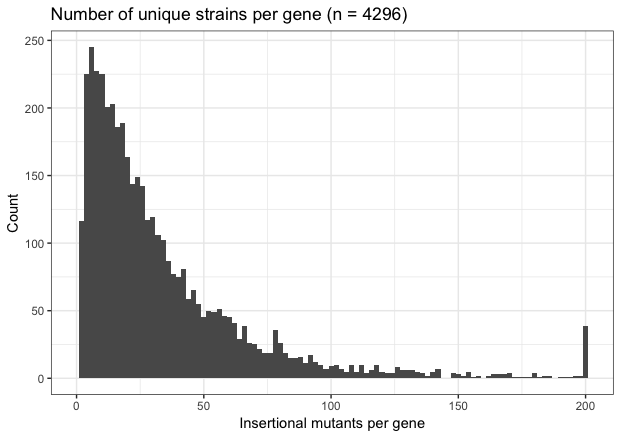

Fig. S2. Of the 920 genes for which their fitness contribution could not be calculated, 408 have no insertion strains in the library, while 512 have at least one insertion but an insufficient number of sequencing reads to calculate fitness. These 920 genes range in size from 73 bp to 4199 bp, with a median size of 575 bp. 7 genes have no TA sites. The histogram of all genes excludes 7 genes containing 139 to 578 TA sites.

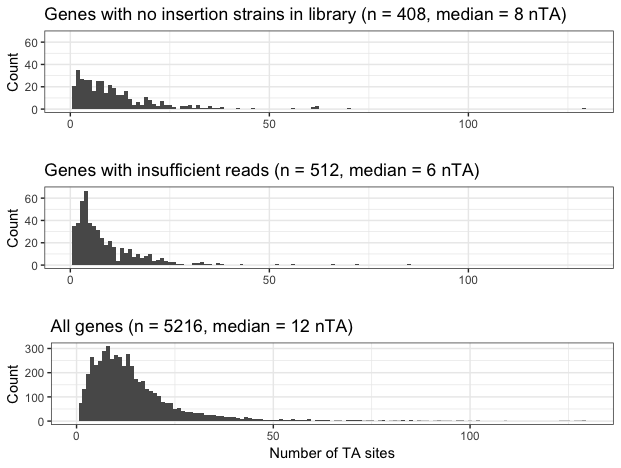

Fig. S3. Representative growth of B728a in the apoplast. B728a WT cells were inoculated into the apoplast by submerging the plants in 1.5 L of inoculum containing 10^5^ CFU/ml in 1 mM KPO_4_ buffer and subjected to a vacuum for 1.25 minutes. Rapidly restoring atmospheric pressure forced the bacterial suspension into the intercellular spaces of the leaves. The plants were allowed to dry on a laboratory bench for at least five hours, and then moved to the greenhouse. Cells were recovered from 8 to 12 leaves at each sampling time by homogenization of the leaves, dilution plating of appropriate dilutions of the homogenate on selective media, followed by enumeration of colonies.

B728a growth in the apoplast

| days post inoculation | log(CFU/g) |
| --- | --- |
| 0 | 4.79742 |
| 4 | 8.55538 |
| 6 | 8.69544 |

Fig. S4. Venn diagrams showing consistency in identifying genes contributing to epiphytic growth (A) and apoplastic growth (B) in different replicate experiments with fitness contribution thresholds of < -2 (left) or < -1 (right), and *t* < -3.

A.

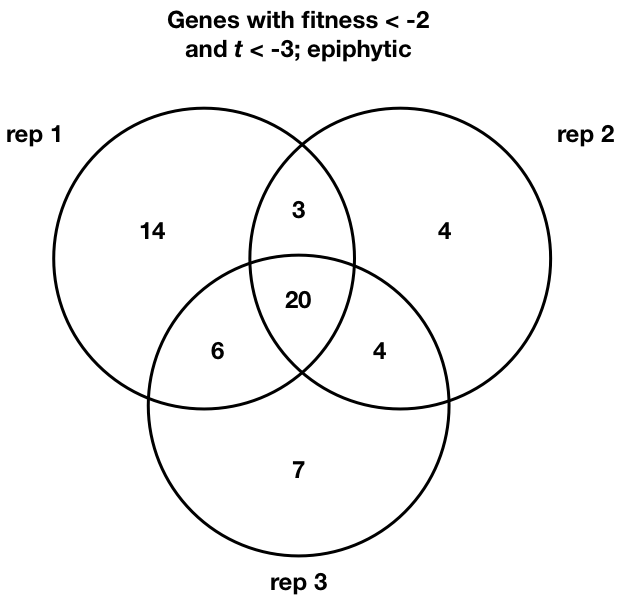

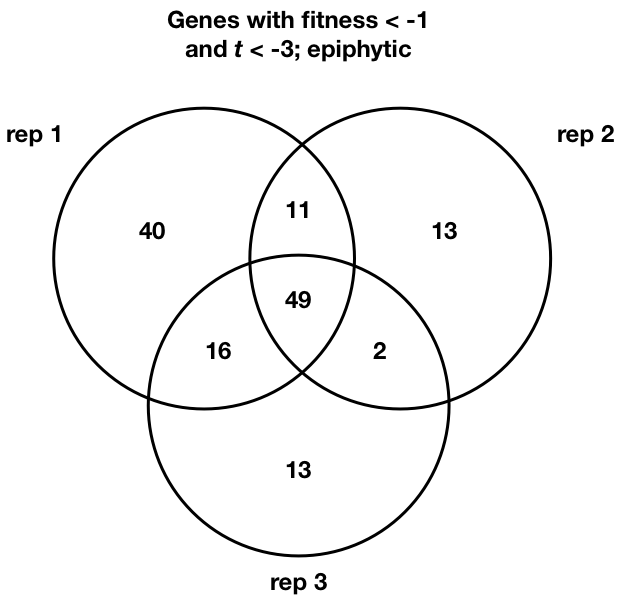

B.

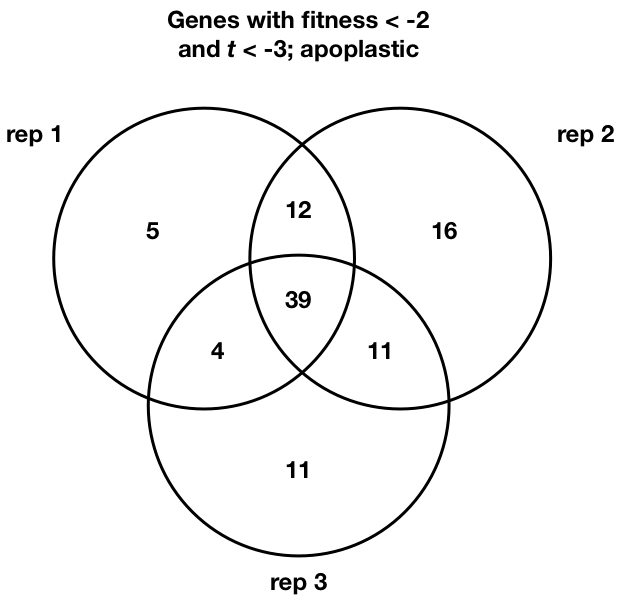

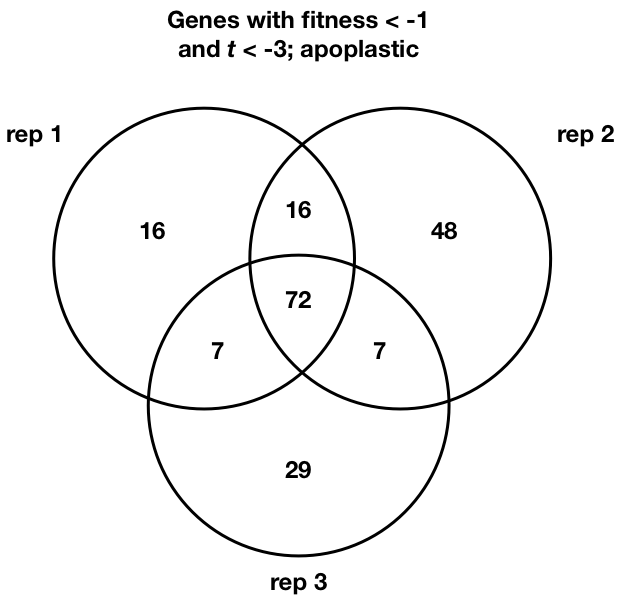

Fig. S5. Scatterplot of average apoplastic fitness and absolute average t-value for all genes, averaging values from three replicate experiments. Genes encoding “Type III secreted proteins” are shown in red, with those encoding “Type III pilus” components and regulatory elements shown in blue. Of the secreted effector proteins, HopAB1 has a consistent but relatively small contribution to apoplastic fitness. Disruptions in individual type III secretion system pilus components have large effects. All other genes are shown in grey.

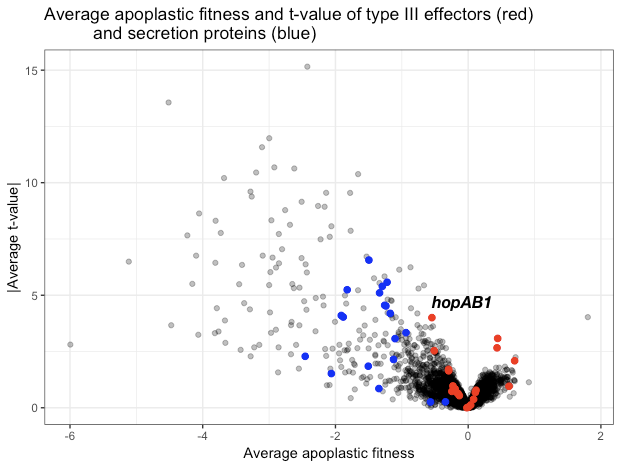

Table S1. Number of genes within each functional category predicted to be essential (N = 392). Functional category annotations for B728a genes are primarily based on COG (7) and KEGG (8) annotations, with manual additions and corrections originally published by Yu *et al.* (9).

| Category | Predicted essential genes |
| --- | --- |
| Translation | 66 |
| None | 38 |
| Cofactor metabolism | 36 |
| Energy generation | 30 |
| Amino acid metabolism and transport | 22 |
| Nucleotide metabolism and transport | 22 |
| Replication and DNA repair | 20 |
| LPS synthesis and transport | 19 |
| Hypothetical | 15 |
| Secretion/Efflux/Export | 15 |
| Peptidoglycan/cell wall polymers | 12 |
| Fatty acid metabolism | 10 |
| Sulfur metabolism and transport | 8 |
| Terpenoid backbone synthesis | 8 |
| Carbohydrate metabolism and transport | 7 |
| Cell division | 7 |
| Siderophore synthesis and transport | 7 |
| Transcription | 5 |
| Chaperones/Heat shock proteins | 4 |
| Nitrogen metabolism | 4 |
| Phospholipid metabolism | 4 |
| Transcription - Sigma factor | 4 |
| Transport (organic compounds) | 3 |
| Iron metabolism and transport | 2 |
| Iron-sulfur proteins | 2 |
| Organic acid metabolism and transport | 2 |
| Outer membrane proteins | 2 |
| Oxidative stress tolerance | 2 |
| Post-translational modification | 2 |
| Stress resistance | 2 |
| Transcriptional regulation | 2 |
| Compatible solute synthesis | 1 |
| Osmosensing & regulation | 1 |
| Oxidative stress tolerance (Antioxidant enzyme) | 1 |
| Phage & IS elements | 1 |
| Phosphate metabolism and transport | 1 |
| Polysaccharide synthesis and regulation | 1 |
| Proteases | 1 |
| Signal transduction mechanisms | 1 |
| Transport | 1 |
| Transport (inorganic ions) | 1 |

Table S2. B728a genes predicted to be essential compared to their predicted homologs in *P. aeruginosa* PAO1. PAO1 homologs to B728a genes were identified using the IMG database genome-gene best homologs function (7). Essential vs. nonessential characterization of PAO1 genes was predicted in (10).

|  | B728a essential | B728a nonessential |
| --- | --- | --- |
| PAO1 essential | 259 | 48 |
| PAO1 nonessential | 104 | 1938 |
| No homolog in PAO1 | 29 | 2610 |
| Total | 392 | 4596 |

Table S3. Genes that when disrupted confer individually large decreases in fitness (average fitness < -2 from two replicate experiments) in King’s B medium. All genes represent those in which *t* < -3 in both replicate experiments. The pooled library was initially generated in LB but inoculum for the experiments was grown on KB. While these are both rich media, the presence of yeast extract in LB apparently provides resources not found in KB and thus the growth of certain biosynthetic mutants was limited in KB. For example, the ability to synthesize the cofactor biotin was required for growth in KB, suggesting its limitation compared to other components such as amino acids and carbon compounds in this medium. However, the presence of biotin auxotrophs in the library indicates that these mutants were able to grow in LB.

| Locus | Name | Description | Fitness in King’s B | | | Classification |
| --- | --- | --- | --- | --- | --- | --- |
|  |  |  | 1 | 2 | Average |  |
| Psyr_4581 | trpG | anthranilate synthase, component II | -2.47 | -1.71 | -2.09 | Amino acid metabolism and transport |
| Psyr_0826 | pgi | glucose-6-phosphate isomerase | -1.91 | -2.21 | -2.06 | Carbohydrate metabolism and transport |
| Psyr_1613 | minC | septum site-determining protein MinC | -3.40 | -3.70 | -3.55 | Cell division |
| Psyr_0454 | bioA | adenosylmethionine-8-amino-7-oxononanoate aminotransferase apoenzyme | -2.56 | -2.62 | -2.59 | Cofactor metabolism |
| Psyr_4687 | bioB | biotin synthase | -2.87 | -2.96 | -2.91 | Cofactor metabolism |
| Psyr_4683 | bioD | dethiobiotin synthase | -3.30 | -2.53 | -2.92 | Cofactor metabolism |
| Psyr_4686 | bioF | 8-amino-7-oxononanoate synthase | -2.89 | -3.18 | -3.03 | Cofactor metabolism |
| Psyr_0951 | moeB | UBA/THIF-type NAD/FAD binding fold:MoeZ/MoeB | -4.81 | -6.07 | -5.44 | Cofactor metabolism |
| Psyr_2080 | pabB | aminodeoxychorismate synthase, subunit I | -1.38 | -2.65 | -2.01 | Cofactor metabolism |
| Psyr_4341 | thiE | thiamine-phosphate diphosphorylase | -2.80 | -3.71 | -3.26 | Cofactor metabolism |
| Psyr_4740 | thiG | thiazole-phosphate synthase | -3.69 | -3.98 | -3.84 | Cofactor metabolism |
| Psyr_4340 |  | phosphomethylpyrimidine kinase, putative | -3.51 | -4.07 | -3.79 | Cofactor metabolism |
| Psyr_0565 |  | Protein of unknown function UPF0126 | -2.35 | -3.33 | -2.84 | Hypothetical |
| Psyr_0917 | rfbA-2 | ABC-2 | -1.90 | -2.53 | -2.21 | LPS synthesis and transport |
| Psyr_0918 | rfbB-2 | ABC transporter | -2.05 | -2.37 | -2.21 | LPS synthesis and transport |
| Psyr_0259 | envZ | Osmolarity sensor protein envZ | -2.29 | -1.91 | -2.10 | Osmosensing & regulation |
| Psyr_3008 | uppP | Undecaprenyl-diphosphatase | -2.95 | -4.69 | -3.82 | Peptidoglycan/cell wall polymers |
| Psyr_0849 | pssA-1 | CDP-diacylglycerol--serine O-phosphatidyltransferase | -2.04 | -2.32 | -2.18 | Phospholipid metabolism |
| Psyr_4091 |  | 8-oxo-dGTPase | -2.54 | -2.47 | -2.51 | Replication and DNA repair |
| Psyr_1544 |  | SirA-like protein | -3.94 | -4.44 | -4.19 |  |

Table S4. Numbers of unique barcodes and median reads per gene in each of three replicate experiments and samples obtained following library outgrowth (time0) and after growth in a particular habitat. Unique barcodes were calculated as the total that mapped to the genome and had 3 or more reads in a given experiment. The total number of mapped barcodes in the library = 281,417. Technical (sequencing) replicates are listed separately (“a” and “b”), and share the same time0 reference sample. For an experiment to pass quality control, the median reads per gene in the sample must be ≥ 50 (2).

| Experiment | Unique barcodes at time0 | Unique barcodes in sample | % Recovery | Median reads/gene at time0 | Median reads/gene in sample |
| --- | --- | --- | --- | --- | --- |
| KB_1 | 187,482 | 192,078 | >100* | 278 | 150 |
| KB_2 | 197,089 | 226,857 | >100* | 378.5 | 251 |
| Epiphytic_1a | 196,894 | 191,648 | 97.3 | 279 | 167 |
| Epiphytic_1b | 196,894 | 182,055 | 92.5 | 279 | 148 |
| Epiphytic_2a | 216,700 | 177,679 | 82.0 | 381 | 175.5 |
| Epiphytic_2b | 216,700 | 173,567 | 80.1 | 381 | 164.5 |
| Epiphytic_3a | 230,482 | 201,171 | 87.3 | 453.5 | 248.5 |
| Epiphytic_3b | 230,482 | 196,807 | 85.4 | 453.5 | 211 |
| Apoplast_1a | 218,149 | 149,311 | 68.4 | 409.5 | 163 |
| Apoplast_1b | 218,149 | 151,211 | 69.3 | 409.5 | 155 |
| Apoplast_2 | 214,346 | 156,416 | 73.0 | 390 | 160 |
| Apoplast_3a | 222,473 | 185,183 | 83.2 | 397 | 203 |
| Apoplast_3b | 222,473 | 187,289 | 84.2 | 397 | 216 |

* More unique barcodes sequenced at the end of an experiment indicates that additional unique barcodes were present but were not sequenced at the start of the experiment (time0).

Table S5. Genes with average fitness < -2 in epiphytic experiments. All genes represent those in which *t* < -3 in at least two experiments.

| Locus | Name | Description | Epiphytic Fitness | | | | Classification |
| --- | --- | --- | --- | --- | --- | --- | --- |
|  |  |  | 1 | 2 | 3 | Average |  |
| Psyr_4270 | glyA | serine hydroxymethyltransferase | -2.35 | -2.94 | -3.42 | -2.90 | Amino acid metabolism and transport |
| Psyr_4369 | proA | glutamate-5-semialdehyde dehydrogenase | -3.02 | -2.42 | -2.84 | -2.76 | Amino acid metabolism and transport |
| Psyr_0704 | proB | glutamate 5-kinase | -2.91 | -1.48 | -2.12 | -2.17 | Amino acid metabolism and transport |
| Psyr_0557 | serB | phosphoserine phosphatase | -2.16 | -4.21 | -3.27 | -3.21 | Amino acid metabolism and transport |
| Psyr_0033 | trpA | tryptophan synthase, alpha chain | -3.26 | -3.21 | -4.03 | -3.50 | Amino acid metabolism and transport |
| Psyr_0034 | trpB | tryptophan synthase, beta chain | -2.95 | -3.04 | -4.01 | -3.33 | Amino acid metabolism and transport |
| Psyr_4609 | trpE | anthranilate synthase, component I | -3.60 | -3.31 | -3.96 | -3.62 | Amino acid metabolism and transport |
| Psyr_1663 | trpF | phosphoribosylanthranilate isomerase | -3.54 | -3.50 | -4.59 | -3.88 | Amino acid metabolism and transport |
| Psyr_4852 |  | D-3-phosphoglycerate dehydrogenase | -2.97 | -3.51 | -3.31 | -3.26 | Amino acid metabolism and transport |
| Psyr_2980 | galU | UDP-glucose pyrophosphorylase | -2.64 | -1.34 | -2.07 | -2.02 | Carbohydrate metabolism and transport |
| Psyr_1613* | minC | septum site-determining protein MinC | -1.46 | -2.27 | -2.53 | -2.09 | Cell division |
| Psyr_0454* | bioA | adenosylmethionine-8-amino-7-oxononanoate aminotransferase apoenzyme | -2.43 | -2.01 | -2.35 | -2.26 | Cofactor metabolism |
| Psyr_4687* | bioB | biotin synthase | -2.03 | -2.76 | -2.05 | -2.28 | Cofactor metabolism |
| Psyr_4686* | bioF | 8-amino-7-oxononanoate synthase | -1.94 | -2.28 | -2.18 | -2.13 | Cofactor metabolism |
| Psyr_3174 | cysG | uroporphyrinogen-III C-methyltransferase / precorrin-2 dehydrogenase | -1.33 | -2.70 | -3.07 | -2.37 | Cofactor metabolism |
| Psyr_0846 | ilvI | acetolactate synthase, large subunit | -2.34 | -2.45 | -2.35 | -2.38 | Cofactor metabolism |
| Psyr_0827 | panC | pantothenate synthetase | -2.60 | -4.00 | -2.69 | -3.10 | Cofactor metabolism |
| Psyr_0532 |  | conserved hypothetical protein | -3.40 | -2.35 | -1.73 | -2.50 | Hypothetical |
| Psyr_2461 |  | Uncharacterized conserved protein UCP030820 | -2.13 | -1.79 | -2.44 | -2.12 | Hypothetical |
| Psyr_0917* | rfbA-2 | ABC-2 | -3.05 | -2.13 | -2.62 | -2.60 | LPS synthesis and transport |
| Psyr_0918* | rfbB-2 | ABC transporter | -2.83 | -2.00 | -2.05 | -2.29 | LPS synthesis and transport |
| Psyr_0411 | gltB | glutamate synthase (NADPH) large subunit | -2.73 | -1.65 | -1.64 | -2.01 | Nitrogen metabolism |
| Psyr_1668 | purF | amidophosphoribosyltransferase | -3.20 | -3.21 | -3.15 | -3.18 | Nucleotide metabolism and transport |
| Psyr_1269 | purL | phosphoribosylformylglycinamidine synthase | -3.08 | -2.52 | -2.54 | -2.71 | Nucleotide metabolism and transport |
| Psyr_3008* | uppP | Undecaprenyl-diphosphatase | -1.86 | -2.29 | -3.01 | -2.38 | Peptidoglycan/cell wall polymers |
| Psyr_4512 |  | putative phage-related protein | -2.24 | -1.70 | -2.38 | -2.11 | Phage & IS elements |
| Psyr_0377 | mdoG | Periplasmic glucan biosynthesis protein, MdoG | -2.47 | -2.48 | -2.73 | -2.56 | Polysaccharide synthesis and regulation |
| Psyr_0378 | mdoH | Glycosyl transferase, family 2 | -2.38 | -2.37 | -3.05 | -2.60 | Polysaccharide synthesis and regulation |
| Psyr_1408 | ruvC | Holliday junction endonuclease RuvC | -3.43 | -2.74 | -1.71 | -2.63 | Replication and DNA repair |
| Psyr_2462 |  | Nitrite/sulfite reductase, hemoprotein beta-component, ferrodoxin-like:Nitrite and sulphite reductase 4Fe-4S region | -2.60 | -3.62 | -3.36 | -3.19 | Sulfur metabolism and transport |
| Psyr_0529 |  | Glycosyl transferase, group 1 | -3.59 | -3.11 | -2.88 | -3.19 |  |

* These genes also have a fitness of < -2 and *t* < -3 for growth in KB.

Table S6. Genes with average fitness < -2 in apoplastic experiments. All genes represent those in which *t* < -3 in at least two experiments.

| Locus | Name | Description | Apoplastic Fitness | | | | Classification |
| --- | --- | --- | --- | --- | --- | --- | --- |
|  |  |  | 1 | 2 | 3 | Average |  |
| Psyr_4270 | glyA | serine hydroxymethyltransferase | -5.00 | -5.34 | -5.01 | -5.11 | Amino acid metabolism and transport |
| Psyr_4897 | hisB | imidazoleglycerol-phosphate dehydratase | -3.76 | -3.42 | -2.69 | -3.29 | Amino acid metabolism and transport |
| Psyr_4132 | hisC | histidinol phosphate aminotransferase apoenzyme | -1.90 | -3.58 | -2.57 | -2.69 | Amino acid metabolism and transport |
| Psyr_4133 | hisD | histidinol dehydrogenase | -2.55 | -3.66 | -2.79 | -3.00 | Amino acid metabolism and transport |
| Psyr_4134 | hisG | ATP phosphoribosyltransferase (homohexameric) | -2.51 | -3.37 | -2.40 | -2.76 | Amino acid metabolism and transport |
| Psyr_4896 | hisH | imidazole glycerol phosphate synthase subunit hisH | -2.02 | -2.56 | -1.96 | -2.18 | Amino acid metabolism and transport |
| Psyr_1257 | leuA | 2-isopropylmalate synthase | -2.16 | -2.56 | -1.95 | -2.22 | Amino acid metabolism and transport |
| Psyr_1985 | leuB | 3-isopropylmalate dehydrogenase | -4.35 | -3.99 | -3.82 | -4.05 | Amino acid metabolism and transport |
| Psyr_1983 | leuC | 3-isopropylmalate dehydratase, large subunit | -4.30 | -4.82 | -3.35 | -4.16 | Amino acid metabolism and transport |
| Psyr_0473 | metW | Methionine biosynthesis MetW | -4.66 | -4.25 | -3.38 | -4.10 | Amino acid metabolism and transport |
| Psyr_0474 | metX | homoserine O-acetyltransferase | -4.04 | -3.02 | -3.97 | -3.68 | Amino acid metabolism and transport |
| Psyr_1669 | metZ | O-succinylhomoserine sulfhydrylase | -4.14 | -4.63 | -3.93 | -4.23 | Amino acid metabolism and transport |
| Psyr_4369 | proA | glutamate-5-semialdehyde dehydrogenase | -3.72 | -2.12 | -3.15 | -3.00 | Amino acid metabolism and transport |
| Psyr_0704 | proB | glutamate 5-kinase | -3.24 | -3.19 | -1.98 | -2.80 | Amino acid metabolism and transport |
| Psyr_0557 | serB | phosphoserine phosphatase | -2.88 | -2.37 | -1.64 | -2.30 | Amino acid metabolism and transport |
| Psyr_0033 | trpA | tryptophan synthase, alpha chain | -2.89 | -3.23 | -1.87 | -2.66 | Amino acid metabolism and transport |
| Psyr_0034 | trpB | tryptophan synthase, beta chain | -4.03 | -3.75 | -0.24 | -2.67 | Amino acid metabolism and transport |
| Psyr_4580 | trpD | anthranilate phosphoribosyltransferase | -4.47 | -4.70 | -4.26 | -4.47 | Amino acid metabolism and transport |
| Psyr_4609 | trpE | anthranilate synthase, component I | -4.87 | -4.58 | -4.10 | -4.51 | Amino acid metabolism and transport |
| Psyr_1663 | trpF | phosphoribosylanthranilate isomerase | -3.43 | -3.93 | -3.64 | -3.67 | Amino acid metabolism and transport |
| Psyr_4852 |  | D-3-phosphoglycerate dehydrogenase | -3.06 | -3.30 | -2.49 | -2.95 | Amino acid metabolism and transport |
| Psyr_4894 |  | 1-(5-phosphoribosyl)-5-[(5- phosphoribosylamino)methylideneamino] imidazole-4-carboxamide isomerase | -1.87 | -3.00 | -2.76 | -2.54 | Amino acid metabolism and transport |
| Psyr_2980 | galU | UDP-glucose pyrophosphorylase | -2.10 | -2.53 | -2.28 | -2.30 | Carbohydrate metabolism and transport |
| Psyr_0826* | pgi | glucose-6-phosphate isomerase | -2.87 | -2.96 | -3.46 | -3.10 | Carbohydrate metabolism and transport |
| Psyr_3179 | ftsK | DNA translocase FtsK | -2.86 | -2.53 | -2.23 | -2.54 | Cell division |
| Psyr_1613* | minC | septum site-determining protein MinC | -3.94 | -3.75 | -3.68 | -3.79 | Cell division |
| Psyr_4194 | dnaJ-1 | DnaJ central region:Heat shock protein DnaJ, N-terminal:Chaperone DnaJ, C-terminal | -1.85 | -2.30 | -1.87 | -2.01 | Chaperones/Heat shock proteins |
| Psyr_0454* | bioA | adenosylmethionine-8-amino-7-oxononanoate aminotransferase apoenzyme | -2.32 | -2.17 | -2.07 | -2.19 | Cofactor metabolism |
| Psyr_4687* | bioB | biotin synthase | -2.83 | -2.85 | -2.88 | -2.85 | Cofactor metabolism |
| Psyr_4683* | bioD | dethiobiotin synthase | -2.99 | -2.49 | -3.21 | -2.90 | Cofactor metabolism |
| Psyr_4686* | bioF | 8-amino-7-oxononanoate synthase | -2.54 | -2.81 | -2.53 | -2.63 | Cofactor metabolism |
| Psyr_0848 | ilvC | ketol-acid reductoisomerase | -2.91 | -3.88 | -2.11 | -2.97 | Cofactor metabolism |
| Psyr_0469 | ilvD | dihydroxyacid dehydratase | -3.70 | -4.02 | -2.11 | -3.28 | Cofactor metabolism |
| Psyr_0847 | ilvH | acetolactate synthase, small subunit | -2.85 | -2.72 | -1.72 | -2.43 | Cofactor metabolism |
| Psyr_0846 | ilvI | acetolactate synthase, large subunit | -3.29 | -3.70 | -2.33 | -3.11 | Cofactor metabolism |
| Psyr_4341* | thiE | thiamine-phosphate diphosphorylase | -1.94 | -2.69 | -2.91 | -2.51 | Cofactor metabolism |
| Psyr_4740* | thiG | thiazole-phosphate synthase | -1.93 | -2.18 | -3.84 | -2.65 | Cofactor metabolism |
| Psyr_4340* |  | phosphomethylpyrimidine kinase, putative | -2.01 | -2.79 | -3.76 | -2.85 | Cofactor metabolism |
| Psyr_0167 |  | hypothetical protein | -2.86 | -3.64 | -2.41 | -2.97 | Hypothetical |
| Psyr_1614 | htrB | lipid A biosynthesis acyltransferase | -4.30 | -4.58 | -2.57 | -3.82 | LPS synthesis and transport |
| Psyr_0917* | rfbA-2 | ABC-2 | -2.43 | -3.24 | -3.27 | -2.98 | LPS synthesis and transport |
| Psyr_0918* | rfbB-2 | ABC transporter | -2.08 | -2.40 | -3.38 | -2.62 | LPS synthesis and transport |
| Psyr_0014 |  | lipid A biosynthesis acyltransferase | -3.14 | -2.06 | -2.81 | -2.67 | LPS synthesis and transport |
| Psyr_1668 | purF | amidophosphoribosyltransferase | -3.64 | -3.27 | -4.53 | -3.81 | Nucleotide metabolism and transport |
| Psyr_1269 | purL | phosphoribosylformylglycinamidine synthase | -3.19 | -4.03 | -4.07 | -3.76 | Nucleotide metabolism and transport |
| Psyr_3008* | uppP | Undecaprenyl-diphosphatase | -3.42 | -3.44 | -3.36 | -3.41 | Peptidoglycan/cell wall polymers |
| Psyr_2613 | syrD | Cyclic peptide transporter | -2.77 | -2.88 | -1.70 | -2.45 | Phytotoxin synthesis and transport |
| Psyr_0219 | algC | phosphomannomutase | -3.35 | -2.67 | -3.77 | -3.26 | Polysaccharide synthesis and regulation |
| Psyr_1056 | algL | Poly(beta-D-mannuronate) lyase | -3.02 | -2.82 | -4.29 | -3.38 | Polysaccharide synthesis and regulation |
| Psyr_0377 | mdoG | Periplasmic glucan biosynthesis protein, MdoG | -3.27 | -4.42 | -3.49 | -3.73 | Polysaccharide synthesis and regulation |
| Psyr_0378 | mdoH | Glycosyl transferase, family 2 | -3.83 | -3.70 | -3.89 | -3.81 | Polysaccharide synthesis and regulation |
| Psyr_1350 | mucP | site-2 protease, Metallo peptidase, MEROPS family M50B | -2.71 | -3.14 | -3.11 | -2.99 | Polysaccharide synthesis and regulation |
| Psyr_1748 | clpX | ATP-dependent Clp protease ATP-binding subunit ClpX | -2.97 | -2.42 | -3.29 | -2.90 | Proteases |
| Psyr_1410 | ruvB | Holliday junction DNA helicase RuvB | -1.88 | -2.79 | -2.74 | -2.47 | Replication and DNA repair |
| Psyr_0919 |  | Chromosome segregation ATPase-like protein | -2.02 | -1.91 | -2.49 | -2.14 | Replication and DNA repair |
| Psyr_3958 | algU | RNA polymerase, sigma-24 subunit, RpoE | -2.78 | -2.67 | -3.31 | -2.92 | Transcription - Sigma factor |
| Psyr_4408 | retS | Response regulator receiver:ATP-binding region, ATPase-like:Histidine kinase A, N-terminal:Histidine kinase:Histidine kinase | -2.50 | -2.32 | -2.71 | -2.51 | Transcriptional regulation |
| Psyr_3637 | wbpL | Glycosyl transferase, family 4 | -2.50 | -2.04 | -1.95 | -2.16 |  |
| Psyr_0936 | wbpY | Glycosyl transferase, group 1 | -1.90 | -1.68 | -2.61 | -2.06 |  |
| Psyr_0914 | wbpZ | Glycosyl transferase, group 1 | -2.22 | -1.78 | -2.26 | -2.09 |  |
| Psyr_0529 |  | Glycosyl transferase, group 1 | -3.78 | -2.89 | -3.69 | -3.45 |  |
| Psyr_0915 |  | NAD-dependent epimerase/dehydratase | -3.09 | -3.05 | -3.44 | -3.20 |  |
| Psyr_1544* |  | SirA-like protein | -1.67 | -2.64 | -3.01 | -2.44 |  |
| Psyr_0920 |  | Glycosyl transferase, group 1 | -2.69 | -2.34 | -2.24 | -2.42 |  |
| Psyr_4130 |  | Peptidase S1, chymotrypsin:PDZ/DHR/GLGF | -2.51 | -2.20 | -2.08 | -2.26 |  |

* These genes also fulfill average fitness < -2 and *t* < -3 for growth in KB.

Regarding *wbpYZ*: in the original gene metadata these genes are annotated as “*wpbY*” and “*wpbZ*”. This is likely a typo, as *wbp* genes are involved in O-antigen biosynthesis in *P. aeruginosa*, and “*wpb*” genes could not be identified in *P. syringae* or other Pseudomonads.

Table S7. Functional categories that are enriched among genes important for fitness *in planta*. For this analysis, we only included genes with average fitness < -2 in the *in planta* habitats. The KB data are shown for comparison.

Hypergeometic test for enrichment, p-values across categories:

| **Category** | **Category size**  **(total)** | **King’s B** | **Epiphytic** | **Apoplastic** |
| --- | --- | --- | --- | --- |
| Amino acid metabolism and transport | 210 | 0.329 | 3.294E-08 | 5.127E-16 |
| Polysaccharide synthesis and regulation | 49 | 0.241 | 0.011 | 3.332E-04 |
| Nucleotide metabolism and transport | 48 | 0.029 | 7.801E-05 | 2.244E-03 |
| Type III secretion system | 42 | 0.211 | 0.333 | 8.653E-03 |
| Phytotoxin synthesis and transport | 24 | 0.126 | 0.206 | 0.01085 |

Table S8. Genes with average fitness < -1 but > -2 in King’s B (A), epiphytic (B) and apoplastic experiments (C). All genes represent those in which *t* < -3 in at least two experiments.

A.

| Locus | Name | Description | Fitness in King’s B | | | Classification |
| --- | --- | --- | --- | --- | --- | --- |
|  |  |  | 1 | 2 | Average |  |
| Psyr_2855 | metE | methionine synthase (B12-independent) | -1.70 | -1.34 | -1.52 | Amino acid metabolism and transport |
| Psyr_1751 | ppiD | PpiC-type peptidyl-prolyl cis-trans isomerase | -1.44 | -1.17 | -1.31 | Chaperones/Heat shock proteins |
| Psyr_1650 | pabC | aminodeoxychorismate lyase apoprotein | -1.66 | -2.18 | -1.92 | Cofactor metabolism |
| Psyr_0475 |  | Protein of unknown function YGGT | -2.19 | -1.37 | -1.78 | Hypothetical |
| Psyr_3581 |  | ribosomal large subunit pseudouridine synthase B | -1.31 | -1.08 | -1.19 | Nucleotide metabolism and transport |
| Psyr_2854 |  | conserved hypothetical protein | -1.80 | -1.76 | -1.78 | Phage & IS elements |
| Psyr_0377 | mdoG | Periplasmic glucan biosynthesis protein, MdoG | -1.39 | -1.62 | -1.50 | Polysaccharide synthesis and regulation |
| Psyr_0378 | mdoH | Glycosyl transferase, family 2 | -0.81 | -1.27 | -1.04 | Polysaccharide synthesis and regulation |
| Psyr_1748 | clpX | ATP-dependent Clp protease ATP-binding subunit ClpX | -1.09 | -1.55 | -1.32 | Proteases |
| Psyr_0574 | hflK | protease FtsH subunit HflK | -1.51 | -0.81 | -1.16 | Proteases |
| Psyr_1749 | lon-1 | ATP-dependent proteinase, Serine peptidase, MEROPS family S16 | -0.97 | -1.09 | -1.03 | Proteases |
| Psyr_1410 | ruvB | Holliday junction DNA helicase RuvB | -1.35 | -1.33 | -1.34 | Replication and DNA repair |
| Psyr_4408 | retS | Response regulator receiver:ATP-binding region, ATPase-like:Histidine kinase A, N-terminal:Histidine kinase:Histidine kinase | -1.28 | -0.92 | -1.10 | Transcriptional regulation |
| Psyr_3954 | lepA | GTP-binding protein LepA | -1.17 | -1.56 | -1.37 |  |
| Psyr_4424 |  | Propeptide, PepSY amd peptidase M4:PepSY-associated TM helix | -0.91 | -1.20 | -1.05 |  |
| Psyr_0529 |  | Glycosyl transferase, group 1 | -1.57 | -1.48 | -1.52 |  |

B.

| Locus | Name | Description | Epiphytic Fitness | | | | Classification |
| --- | --- | --- | --- | --- | --- | --- | --- |
|  |  |  | 1 | 2 | 3 | Average |  |
| Psyr_4132 | hisC | histidinol phosphate aminotransferase apoenzyme | -1.76 | -1.67 | -1.17 | -1.54 | Amino acid metabolism and transport |
| Psyr_4133 | hisD | histidinol dehydrogenase | -1.87 | -0.89 | -1.03 | -1.27 | Amino acid metabolism and transport |
| Psyr_4134 | hisG | ATP phosphoribosyltransferase (homohexameric) | -1.63 | -1.11 | -0.84 | -1.19 | Amino acid metabolism and transport |
| Psyr_4896 | hisH | imidazole glycerol phosphate synthase subunit hisH | -2.68 | -1.93 | -1.20 | -1.94 | Amino acid metabolism and transport |
| Psyr_1985 | leuB | 3-isopropylmalate dehydrogenase | -1.58 | -1.19 | -0.90 | -1.23 | Amino acid metabolism and transport |
| Psyr_1983 | leuC | 3-isopropylmalate dehydratase, large subunit | -1.53 | -2.03 | -0.41 | -1.33 | Amino acid metabolism and transport |
| Psyr_4894 |  | 1-(5-phosphoribosyl)-5-[(5- phosphoribosylamino)methylideneamino] imidazole-4-carboxamide isomerase | -2.42 | -1.35 | -1.22 | -1.66 | Amino acid metabolism and transport |
| Psyr_0916 | gmd | GDP-mannose 4,6-dehydratase | -1.64 | -0.61 | -0.77 | -1.01 | Carbohydrate metabolism and transport |
| Psyr_0826 | pgi | glucose-6-phosphate isomerase | -2.08 | -1.05 | -2.55 | -1.89 | Carbohydrate metabolism and transport |
| Psyr_4842 |  | Phosphoenolpyruvate-protein phosphotransferase | -1.67 | -1.49 | -1.13 | -1.43 | Carbohydrate metabolism and transport |
| Psyr_3179 | ftsK | DNA translocase FtsK | -1.98 | -1.15 | -1.90 | -1.68 | Cell division |
| Psyr_0848 | ilvC | ketol-acid reductoisomerase | -1.11 | -1.40 | -1.62 | -1.38 | Cofactor metabolism |
| Psyr_0469 | ilvD | dihydroxyacid dehydratase | -1.62 | -2.48 | -1.64 | -1.91 | Cofactor metabolism |
| Psyr_0847 | ilvH | acetolactate synthase, small subunit | -2.09 | -2.01 | -1.90 | -2.00 | Cofactor metabolism |
| Psyr_1120 | zwf-1 | glucose-6-phosphate 1-dehydrogenase | -1.09 | -1.37 | -0.60 | -1.02 | Energy generation |
| Psyr_3691 |  | conserved hypothetical protein | -1.53 | -0.22 | -1.32 | -1.03 | Hypothetical |
| Psyr_0534 |  | membrane protein, putative | -1.70 | -0.76 | -1.11 | -1.19 | Hypothetical |
| Psyr_3805 |  | hypothetical protein | -2.26 | 0.57 | -2.54 | -1.41 | Hypothetical |
| Psyr_0923 |  | hypothetical protein | -1.64 | -1.28 | -1.43 | -1.45 | Hypothetical |
| Psyr_0983 |  | Protein of unknown function DUF159 | -1.93 | -1.43 | -1.64 | -1.67 | Hypothetical |
| Psyr_0412 | gltD | glutamate synthase (NADPH) small subunit | -1.85 | -0.65 | -0.78 | -1.09 | Nitrogen metabolism |
| Psyr_0294 | ppx | Exopolyphosphatase | -0.87 | -0.78 | -1.58 | -1.08 | Nucleotide metabolism and transport |
| Psyr_3581 |  | ribosomal large subunit pseudouridine synthase B | -0.61 | -1.44 | -1.57 | -1.21 | Nucleotide metabolism and transport |
| Psyr_0976 | mqo | Malate:quinone-oxidoreductase | -1.89 | -1.55 | -1.62 | -1.69 | Organic acid metabolism and transport |
| Psyr_0630 | mpl | UDP-N-acetylmuramate:L-alanyl-gamma-D-glutamyl- meso-diaminopimelate ligase | -1.62 | -0.82 | -1.23 | -1.22 | Peptidoglycan/cell wall polymers |
| Psyr_0402 | ponA | Peptidoglycan glycosyltransferase | -1.20 | -0.78 | -1.17 | -1.05 | Peptidoglycan/cell wall polymers |
| Psyr_4158 | eftA | conserved hypothetical protein | -1.85 | -0.52 | -1.32 | -1.23 | Plant-associated proteins |
| Psyr_3636 | wbpM | Polysaccharide biosynthesis protein CapD | -1.56 | -0.76 | -1.07 | -1.13 | Polysaccharide synthesis and regulation |
| Psyr_4887 | ctpA | Peptidase S41A, C-terminal protease | -2.42 | -1.64 | -0.98 | -1.68 | Post-translational modification |
| Psyr_1748 | clpX | ATP-dependent Clp protease ATP-binding subunit ClpX | -1.47 | -1.27 | -1.06 | -1.27 | Proteases |
| Psyr_0201 | recG | ATP-dependent DNA helicase RecG | -1.50 | -0.58 | -0.99 | -1.02 | Replication and DNA repair |
| Psyr_1410 | ruvB | Holliday junction DNA helicase RuvB | -1.65 | -1.38 | -2.38 | -1.80 | Replication and DNA repair |
| Psyr_5065 | uvrD | ATP-dependent DNA helicase UvrD | -1.84 | -1.41 | -2.30 | -1.85 | Replication and DNA repair |
| Psyr_0919 |  | Chromosome segregation ATPase-like protein | -1.66 | -1.05 | -1.16 | -1.29 | Replication and DNA repair |
| Psyr_0832 | cbrA-1 | Two-component sensor kinase CbrA | -1.28 | -1.65 | -1.35 | -1.43 | Signal transduction mechanisms |
| Psyr_0811 | terC | Integral membrane protein TerC | -1.11 | -0.74 | -1.46 | -1.10 | Stress resistance |
| Psyr_4128 | cysD | sulfate adenylyltransferase subunit 2 | -1.28 | -1.07 | -0.83 | -1.06 | Sulfur metabolism and transport |
| Psyr_4126 | cysNC | adenylylsulfate kinase / sulfate adenylyltransferase subunit 1 | -1.08 | -1.15 | -1.40 | -1.21 | Sulfur metabolism and transport |
| Psyr_4408 | retS | Response regulator receiver:ATP-binding region, ATPase-like:Histidine kinase A, N-terminal:Histidine kinase:Histidine kinase | -2.05 | -1.39 | -1.10 | -1.51 | Transcriptional regulation |
| Psyr_4362 | rlpA-2 | Rare lipoprotein A | -2.16 | -1.93 | -1.67 | -1.92 |  |
| Psyr_3637 | wbpL | Glycosyl transferase, family 4 | -1.68 | -1.09 | -1.04 | -1.27 |  |
| Psyr_0936 | wpbY | Glycosyl transferase, group 1 | -1.77 | -0.52 | -0.87 | -1.06 |  |
| Psyr_0914 | wpbZ | Glycosyl transferase, group 1 | -1.65 | -0.78 | -0.83 | -1.09 |  |
| Psyr_4078 |  | AmpG-related permease | -1.32 | -0.74 | -1.02 | -1.03 |  |
| Psyr_4844 |  | HAD-superfamily hydrolase, subfamily IB (PSPase-like):HAD-superfamily subfamily IB hydrolase, hypothetical 2 | -1.60 | -0.97 | -0.66 | -1.08 |  |
| Psyr_1544 |  | SirA-like protein | -1.92 | -1.54 | 0.22 | -1.08 |  |
| Psyr_4623 |  | Aminoglycoside phosphotransferase | -1.56 | -0.96 | -0.83 | -1.12 |  |
| Psyr_0915 |  | NAD-dependent epimerase/dehydratase | -1.63 | -0.97 | -0.92 | -1.17 |  |
| Psyr_0947 |  | TPR repeat protein:TPR repeat protein | -1.54 | -1.02 | -1.08 | -1.21 |  |
| Psyr_0920 |  | Glycosyl transferase, group 1 | -1.78 | -0.95 | -1.05 | -1.26 |  |
| Psyr_4622 |  | Nucleotidyl transferase | -1.65 | -1.47 | -0.93 | -1.35 |  |

C.

| Locus | Name | Description | Apoplastic Fitness | | | | Classification |
| --- | --- | --- | --- | --- | --- | --- | --- |
|  |  |  | 1 | 2 | 3 | Average |  |
| Psyr_0385 | hisE | phosphoribosyl-ATP pyrophosphatase | -2.30 | -2.21 | -0.92 | -1.81 | Amino acid metabolism and transport |
| Psyr_1109 | edd | 6-phosphogluconate dehydratase | -0.95 | -1.86 | -0.69 | -1.17 | Carbohydrate metabolism and transport |
| Psyr_0916 | gmd | GDP-mannose 4,6-dehydratase | -1.06 | -1.38 | -1.58 | -1.34 | Carbohydrate metabolism and transport |
| Psyr_0758 | scrB | beta-fructofuranosidase | -1.44 | -2.02 | -1.52 | -1.66 | Carbohydrate metabolism and transport |
| Psyr_1914 | talB | transaldolase | -0.96 | -1.82 | -0.67 | -1.15 | Carbohydrate metabolism and transport |
| Psyr_3174 | cysG | uroporphyrinogen-III C-methyltransferase / precorrin-2 dehydrogenase | -1.89 | -1.78 | -0.39 | -1.35 | Cofactor metabolism |
| Psyr_1542 | nadA | quinolinate synthetase | -1.65 | -1.15 | -0.67 | -1.16 | Cofactor metabolism |
| Psyr_5130 |  | chromosome segregation ATPase | -0.91 | -1.18 | -1.18 | -1.09 | Cofactor metabolism |
| Psyr_0532 |  | conserved hypothetical protein | -1.83 | -1.16 | -1.68 | -1.56 | Hypothetical |
| Psyr_0534 |  | membrane protein, putative | -1.26 | -2.07 | -2.00 | -1.78 | Hypothetical |
| Psyr_5067 |  | conserved hypothetical protein | -2.92 | -1.07 | -1.53 | -1.84 | Hypothetical |
| Psyr_0923 |  | hypothetical protein | -1.67 | -1.56 | -2.34 | -1.86 | Hypothetical |
| Psyr_3690 | purN | formyltetrahydrofolate-dependent phosphoribosylglycinamide formyltransferase | -1.84 | -1.13 | -1.25 | -1.41 | Nucleotide metabolism and transport |
| Psyr_4018 | purU | Formyltetrahydrofolate deformylase | -1.76 | -1.39 | -1.82 | -1.66 | Nucleotide metabolism and transport |
| Psyr_2601 | salA | regulatory protein, LuxR | -1.46 | -1.14 | -1.05 | -1.21 | Phytotoxin synthesis and transport |
| Psyr_4158 | eftA | conserved hypothetical protein | -1.99 | -0.99 | -1.28 | -1.42 | Plant-associated proteins |
| Psyr_1055 | algI | Membrane bound O-acyl transferase, MBOAT | -1.02 | -1.00 | -1.10 | -1.04 | Polysaccharide synthesis and regulation |
| Psyr_1054 | algJ | alginate biosynthesis protein AlgJ | -1.50 | -1.76 | -0.94 | -1.40 | Polysaccharide synthesis and regulation |
| Psyr_3636 | wbpM | Polysaccharide biosynthesis protein CapD | -1.88 | -1.87 | -1.56 | -1.77 | Polysaccharide synthesis and regulation |
| Psyr_5065 | uvrD | ATP-dependent DNA helicase UvrD | -1.52 | -0.96 | -1.01 | -1.16 | Replication and DNA repair |
| Psyr_4091 |  | 8-oxo-dGTPase | -1.52 | -1.76 | -1.46 | -1.58 | Replication and DNA repair |
| Psyr_0579 | rnr | RNAse R | -1.79 | -1.32 | -0.67 | -1.26 | RNA degradation |
| Psyr_4843 |  | NUDIX hydrolase | -0.87 | -1.97 | -0.99 | -1.28 | RNA degradation |
| Psyr_4882 | secB | protein translocase subunit secB | -1.27 | -0.69 | -1.46 | -1.14 | Secretion/Efflux/Export |
| Psyr_4069 | colS | ATP-binding region, ATPase-like:Histidine kinase, HAMP region:Histidine kinase A, N-terminal | -1.93 | -0.79 | -0.93 | -1.22 | Signal transduction mechanisms |
| Psyr_2077 | cysB | regulatory protein, LysR:LysR, substrate-binding protein | -1.45 | -1.17 | -1.12 | -1.24 | Sulfur metabolism and transport |
| Psyr_5133 | trmE | tRNA modification GTPase trmE | -0.74 | -1.77 | -1.29 | -1.27 | Translation |
| Psyr_1200 | hrcC | outer-membrane type III secretion protein HrcC | -1.76 | -1.15 | -0.76 | -1.22 | Type III secretion system |
| Psyr_1213 | hrcN | type III secretion cytoplasmic ATPase HrcN | -2.08 | -1.78 | -1.61 | -1.82 | Type III secretion system |
| Psyr_1208 | hrcR | type III secretion protein HrcR | -1.93 | -1.56 | -1.00 | -1.50 | Type III secretion system |
| Psyr_1205 | hrcU | type III secretion protein HrcU | -1.61 | -0.99 | -0.92 | -1.17 | Type III secretion system |
| Psyr_1215 | hrcV | Type III secretion protein HrcV | -1.25 | -1.24 | -1.23 | -1.24 | Type III secretion system |
| Psyr_1197 | hrpE | type III secretion protein HrpE | -1.11 | -2.74 | -1.79 | -1.88 | Type III secretion system |
| Psyr_1216 | hrpJ | type III secretion outer membrane protein PopN | -1.58 | -1.48 | -0.94 | -1.34 | Type III secretion system |
| Psyr_1218 | hrpK1 | type III helper protein HrpK1 | -1.22 | -1.50 | -1.15 | -1.29 | Type III secretion system |
| Psyr_1211 | hrpP | type III secretion protein HrpP | -2.00 | -1.87 | -1.87 | -1.91 | Type III secretion system |
| Psyr_1191 | hrpS | type III transcriptional regulator HrpS | -1.62 | -1.21 | -0.94 | -1.26 | Type III secretion system |
| Psyr_4362 | rlpA-2 | Rare lipoprotein A | -2.88 | -0.85 | -2.10 | -1.95 |  |
| Psyr_4844 |  | HAD-superfamily hydrolase, subfamily IB (PSPase-like):HAD-superfamily subfamily IB hydrolase, hypothetical 2 | -0.96 | -1.16 | -0.92 | -1.01 |  |
| Psyr_1417 |  | TPR repeat protein | -1.06 | -1.09 | -0.89 | -1.02 |  |
| Psyr_4566 |  | Peptidase M23B | -2.14 | -1.11 | -1.34 | -1.53 |  |
| Psyr_1395 |  | virulence | -1.40 | -1.50 | -1.84 | -1.58 |  |
